# The mitotic stopwatch synergizes with mild p53 activation to halt cell proliferation

**DOI:** 10.64898/2025.12.02.691971

**Authors:** Beata E. Mierzwa, Franz Meitinger, Arshad Desai, Karen Oegema

**Affiliations:** Department of Cell and Developmental Biology, School of Biological Sciences, University of California, San Diego, La Jolla, CA 92093, USA; Department of Cellular and Molecular Medicine, University of California, San Diego, La Jolla, California 92093, USA; Okinawa Institute of Science and Technology Graduate University, Okinawa 904-0495, Japan

**Keywords:** mitotic stopwatch, mitotic surveillance pathway, p53, USP28, aneuploidy

## Abstract

The mitotic stopwatch suppresses proliferation of cell lineages experiencing prolonged mitosis that are prone to chromosome missegregation and tumorigenesis. It converts extended mitotic duration into heritable USP28–53BP1 complexes that stabilize p53 and accumulate over generations. To identify genes whose knockout activates the stopwatch, we performed a CRISPR/Cas9 screen comparing dropout kinetics of essential-gene gRNAs in cells lacking versus possessing the stopwatch. Two classes of knockouts emerged: one (27/60 top hits) that prolonged mitosis, and another (33/60 top hits) that mildly elevated p53 without significant mitotic defects, indicating that the stopwatch synergizes with mild p53 activation to halt proliferation. Mild p53 elevation lowered the stopwatch complex threshold for daughter cell arrest and slightly prolonged mitosis. Integrated over successive divisions, the cumulative effect of multiple short mitotic extensions triggered stopwatch-dependent arrest. Thus, the mitotic stopwatch endows the p53 network with a durable lineage memory of modest stress, explaining its tumor-suppressive role.

---

Prolonged mitosis is a hallmark of cells experiencing defects in chromosome alignment or segregation^1–5^, errors that generate aneuploid progeny and promote chromosomal rearrangements that serve as intermediates in nearly all cancers^6, 7^. Suppressing tumorigenesis by identifying and restricting the proliferation of potentially aberrant cells represents a central challenge for tissues with ongoing cell division. This function is accomplished, in part, by a mitotic surveillance mechanism termed the mitotic stopwatch^8–13^. The mitotic stopwatch monitors mitotic duration and, in an analog fashion, converts extensions beyond the normal duration into heritable mitotic stopwatch complexes that are transmitted to daughter cells^12^. Stopwatch complexes require the mitotic kinase Polo-like kinase 1 (PLK1) for their formation and consist of p53-binding protein 1 (53BP1) bound to the deubiquitinase USP28^8–12^. Under normal conditions, p53 is continuously ubiquitinated by its antagonist, the E3 ubiquitin ligase MDM2, which leads to its recognition and destruction by the proteosome^14, 15^. Stopwatch complexes bind p53^12^, which is proposed to enable USP28-dependent p53 deubiquitination and stabilization^9–12^. Thus, prolonged mitosis leads to the production of mitotic stopwatch complexes that are inherited by daughter cells and have the potential to elevate p53 levels.

In RPE1 cells, which have an intact p53 pathway, mitoses exceeding ∼90 minutes (normal duration is ∼20 minutes) generate sufficient stopwatch complexes to trigger an immediate p53/p21-dependent arrest in the daughter cells^12^. Individual prolonged mitoses below this threshold allow continued proliferation. Prior work in RPE1 cells has shown that the mitotic stopwatch can also arrest cells after sequential, modestly prolonged mitoses (50-60 minutes), indicating that the stopwatch integrates mitotic extensions over generations^12^. Because p53 is required for stopwatch-dependent arrest, the ability to monitor mitotic extension is lost in p53-mutant cancers. This ability is also lost in many p53-wildtype cancers, some of which carry mutations in USP28 or 53BP1^8, 12^, consistent with their designation as tumor suppressors^16^.

To address the broader relevance of the mitotic stopwatch in tumor suppression, we performed a multigenerational CRISPR/Cas9 screen comparing guide RNA (gRNA) dropout kinetics in RPE1 cells lacking (*USP28Δ*) versus possessing (WT) the stopwatch. Our goal was to identify essential gene knockouts that activate the stopwatch over multiple successive divisions. Of the top 60 hits, 27 knockouts substantially prolonged mitosis, which would be expected to activate the stopwatch over the course of a few divisions. Interestingly, for the other 33 top hits, the primary effect of was to elevate p53/p21 without affecting mitotic duration, indicating that the mitotic stopwatch also halts proliferation following mild p53 activation. As might be anticipated, mild p53 elevation—via MDM2 inhibition or knockout of p53 regulators—lowered the mitotic time threshold for arrest in cells with an intact stopwatch. However, this reduced threshold remained well above the normal duration of mitosis, raising the question of how mild p53 elevation enhances stopwatch-dependent arrest without overt mitotic delay. Analysis of mitotic timing distributions revealed that mild p53 activation predisposes cells to undergo slightly prolonged mitoses which we show is sufficient to trigger the sensitized stopwatch over multiple generations. These findings indicate that stresses causing mild p53 activation shift the distribution of mitotic timing in a way that triggers heritable stopwatch activation, resulting in the progressive accumulation of stopwatch complexes and cell-cycle arrest. This mechanism suggests that the mitotic stopwatch plays a broader-than-anticipated role in linking transient stress responses to long-term proliferative control and likely explains why the mitotic stopwatch exerts potent tumor-suppressive effects.

## Results

### The mitotic stopwatch accounts for the majority of p53-dependent arrest following prolonged mitosis

The centrally important tumor suppressor p53 coordinates cellular responses to a diverse array of stresses. When activated by signals such as hypoxia, DNA damage, or ribosome dysfunction, p53 induces transcription of genes mediating cell cycle arrest or apoptosis^17, 18^. Uetake and Sluder made the pioneering discovery that p53 also monitors mitotic duration: in RPE1 cells, which have an intact p53 pathway, daughter cells arrest if their mothers remain in mitosis for more than ∼100 minutes^13^ (**Fig. 1a**). Similar observations have been made in other p53-wildtype cell lines^12^. To analyze the impact of mother cell mitotic duration on daughter cell proliferation, asynchronous cell cultures treated for 6 hours with the spindle assembly inhibitor monastrol while being imaged live (**Fig. 1b**). Because individual cells enter mitosis at different times during this interval, they experience a range of mitotic durations, which can be correlated with the fate of their daughters. For control RPE1 cells, daughters of mothers with mitoses longer than 100 minutes nearly always arrest, whereas those with mitoses less than 100 minutes continue dividing (**Fig. 1b**). Under basal conditions, p53 levels remain low due to continuous ubiquitination and proteasomal degradation mediated by the E3 ubiquitin ligase MDM2 (^14, 15, 17, 18^; **Fig. 1c**). Two mechanisms have been proposed to explain this arrest. The first is a PLK1-dependent pathway that generates stopwatch complexes composed of 53BP1 and the deubiquitinase USP28 during prolonged mitosis, which bind and deubiquitinate p53 to stabilize it^19, 20^. The second involves gradual reduction in MDM2 levels during extended mitosis, which reduces p53 ubiquitination^21, 22^. To quantify the contribution of the stopwatch to daughter arrest following prolonged mitosis, we performed a meta-analysis of prior datasets (**Fig. 1d, Extended Data Fig. 1**). In control RPE1 cells, >90% of daughters from mitoses longer than 100 minutes arrested, compared to <10% daughters from mitoses less than 100 minutes. Loss of p53 (*TP53sh*) or its effector p21 (*CDKN1AΔ*) abolished arrest regardless of mitotic duration, confirming that this response is p53-dependent (**Fig. 1d**). Disruption of the stopwatch—via deletion of *USP28* or *TP53BP1*, PLK1 inhibition, or introduction of a mutation into 53BP1 that prevents USP28 binding (G1560K; ^23^)—reduced daughter cell arrest following mitoses longer than 100 minutes from >90% to ∼20%, and eliminated arrest after mitoses less than 100 minutes (**Fig. 1d**). These data indicate that the PLK1-dependent formation of USP28 and 53BP1-containing stopwatch complexes during prolonged mitosis, which are inherited by daughter cells leading to p53 stabilization in the subsequent G1 (**Fig. 1e**), accounts for the majority of p53-dependent arrest following prolonged mitosis, with a potential minor contribution from reduced MDM2 levels.

**Figure 1:**
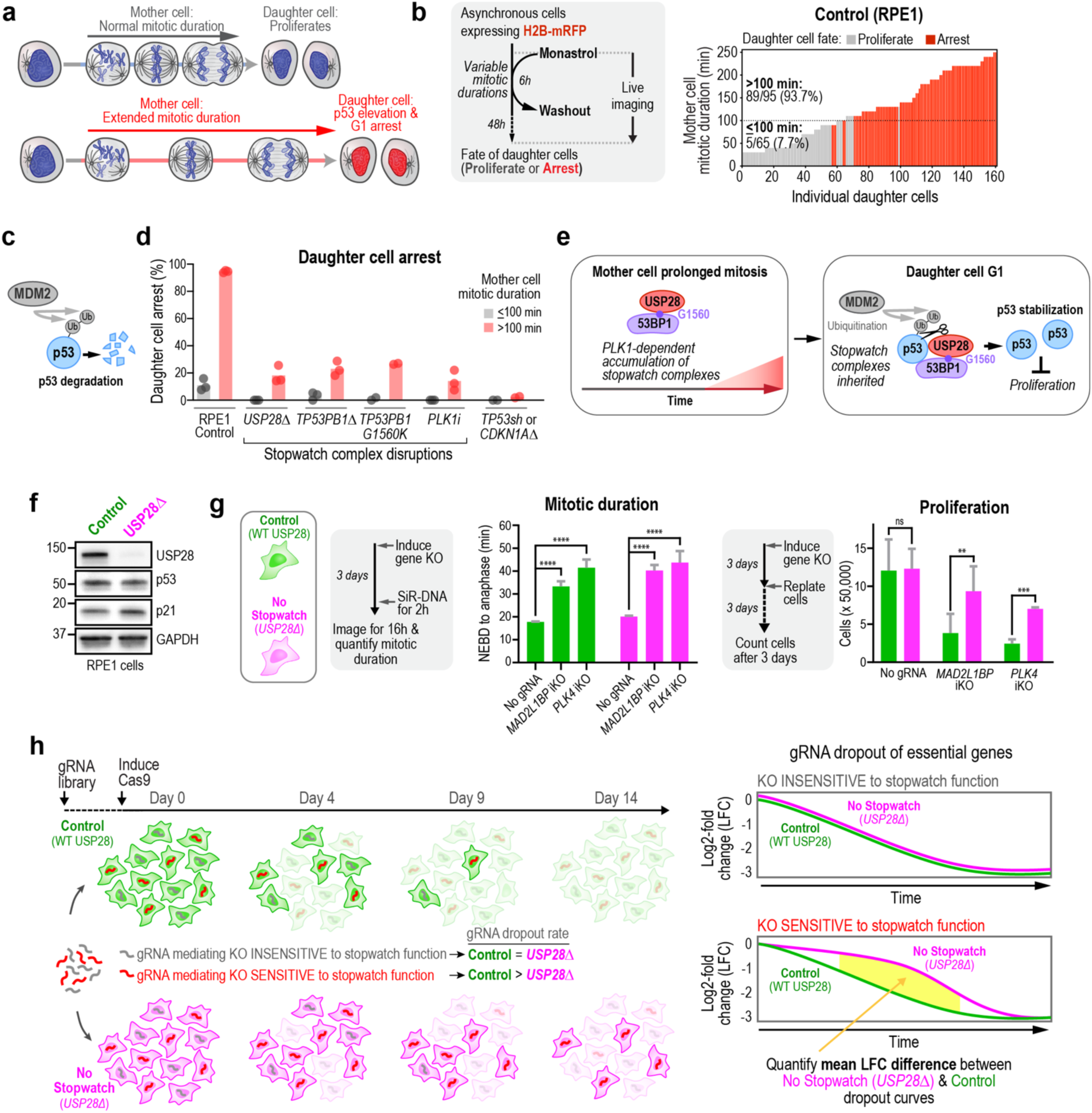
A CRISPR/Cas9 dropout screen to identify essential genes whose knockout makes them sensitive to mitotic stopwatch function. **a,** The p53 pathway monitors mitotic duration: daughters of mother cells that undergo normal mitosis proliferate, whereas daughters of mothers experiencing extended mitosis undergo a p53-dependent G1 arrest. **b**, *Left:* Schematic outlining the stopwatch assay used to assess the relationship between mother cell mitotic duration and daughter cell fate. Asynchronous cells are treated with the spindle-assembly inhibitor monastrol for 6 h to generate mitoses (NEBD to chromosome decondensation) of varying length; daughter cell fate is monitored after washout. *Right:* Stopwatch assay in control RPE1 cells expressing H2B-mRFP. Each bar represents one daughter cell; bar height denotes the mother’s mitotic duration and color indicates whether the daughter divided (*grey*) or arrested (*red*) within 48 h. Percentages of daughter cell arrest for mother cell mitotic durations > or ≤ 100 min are indicated. **c**, Schematic highlights constitutive ubiquitination of p53 by the ubiquitin ligase MDM2 which targets it for proteasomal degradation. **d**, Collated results from 16 stopwatch assays in RPE1 cells, 4 from this study and 12 from prior work^12^. Bars show the mean percentage of daughter cell arrest for mother cell mitotic durations > (*grey*) or ≤ (*red*) 100 min. Points represent independent experiments performed with the same cell line (control), independent clones (*USP28Δ*, *TP53BP1Δ*, *TB53BP1-G1560K*), different inhibitors (*PLK1i*), or knockdown or knockout of different pathway components (*TP53sh*, *CDKN1AΔ*). See **Extended Data Figure 1a** for details. **e**, During prolonged mother cell mitosis, there is a PLK1 kinase-dependent accumulation of USP28–53BP1 stopwatch complexes; the 53BP1-G1560 residue is required for complex formation. In daughter cells, inherited stopwatch complexes deubiquitinate and stabilize p53 to inhibit proliferation. **f,** Immunoblot of USP28, p53, and p21 in control and *USP28Δ* RPE1 cell lines with inducible Cas9 used for the CRISPR screen (see **Extended Data Fig. 1b,c, and Supplementary Table 1**). GAPDH is a loading control. **g,** Experimental schematics (*grey boxes*), and quantified mitotic duration (*middle*) and proliferation (*right*) for control (*green*) or *USP28Δ* (*magenta*) cell lines for the indicated conditions. Error bars represent 95% confidence intervals. Statistical significance was assessed by a two-tailed Welch t-test (**** = P ≤ 0.0001, *** = P ≤ 0.001, ** = P ≤ 0.01, ns = P > 0.05). For mitotic duration, n=124-1315 cells per condition from 2-4 independent experiments; for proliferation, n=3 replicates per condition. **h,** *Left:* Schematic of the CRISPR/Cas9 dropout screen to identify essential genes whose knockout is sensitive to stopwatch function. gRNAs targeting essential genes unrelated to the stopwatch drop out with similar kinetics in the control (*green*) and *USP28Δ* (*magenta*) cell lines, whereas gRNAs targeting essential genes whose knockout is sensitive to mitotic stopwatch function drop out more rapidly in the control compared to the *USP28Δ* cell line. *Right:* The mean difference in log2-fold change (LFC) across the 4 gRNAs per gene between the *USP28Δ* and control cell lines (*yellow shaded region*) quantifies sensitivity of the knockout to stopwatch function.

### A CRISPR/Cas9 screen for essential gene knockouts sensitive to the mitotic stopwatch

To identify essential gene knockouts whose effect on proliferation is impacted by the mitotic stopwatch, we generated matched RPE1 cell lines expressing doxycycline-inducible Cas9 that either possessed (Control) or lacked (*USP28Δ*) the mitotic stopwatch (**Fig. 1f; Extended Data Fig. 1,c;** for a list of all cell lines see **Supplementary Table 1**). To validate these cell lines for screening, we analyzed their proliferation as well as the effects of introducing gRNAs targeting two genes whose knockout prolongs mitosis (**Fig. 1g; Extended Data Fig. 1**). In the absence of any gRNAs, the two cell lines proliferated at equal rates, which supports their use for comparing gRNA dropout kinetics in a screen (**Fig. 1g**). Inducible knockout of PLK4 kinase (*PLK4*), which causes centrosome loss and delayed spindle assembly^24^, or of p31comet (*MAD2L1BP*), which delays spindle checkpoint silencing and thus anaphase onset^12, 25, 26^, resulted in similar increases in mitotic duration—from ∼20 minutes in the ‘no gRNA’ control to ∼40 minutes in cells with either knockout—in both cell lines (**Fig. 1g**). Despite the similar mitotic delays, proliferation was reduced to a significantly greater degree (∼4-fold) in control cells with an intact stopwatch versus the stopwatch-defective *USP28Δ* cells (∼1.5-fold; **Fig. 1g**). These results support the use of these two cell lines to conduct an unbiased screen for gene knockouts whose impact on proliferation is sensitive to mitotic stopwatch function.

We next used the two validated RPE1 cell lines to perform a CRISPR/Cas9 dropout screen with a lentiviral library containing four gRNAs per gene. Control and *USP28Δ* cells were collected at time points between 4 and 14 days after Cas9 induction to monitor gRNA dropout kinetics (**Fig. 1h**; **Extended Data Fig. 1).** gRNAs targeting genes whose knockouts are insensitive to stopwatch function are predicted to drop out with similar kinetics in the two cell lines. In contrast, gRNAs targeting genes whose knockouts are sensitive to the mitotic stopwatch are predicted to drop out faster in the control cell line compared to the *USP28Δ* cell line (**Fig. 1h**, *green vs. magenta*). The screen was performed twice (**Extended Data Fig. 1**) and log_2_-fold change (LFC) values relative to the library plasmid pool were plotted over time for individual gRNAs and for the mean of all four gRNAs per gene (**Fig. 2a**; **Extended Data Fig. 1;** complete dataset available at the Harvard Dataverse repository, https://doi.org/10.7910/DVN/7SFQZ8). For each gene, the LFC was averaged across all time points and then across the two screens to generate a ‘Mean LFC’ per gene in each cell line. The mean LFC in the control cell line was then subtracted from the mean LFC in the *USP28Δ* cell line to yield a single value capturing the difference in dropout rates between *USP28Δ* (*magenta*) and the control (*green*) curves (*yellow area in* **Fig. 1h**, **Fig. 2a**) during the time course. This gap, which we refer to as the ‘Mean LFC Difference’ (*yellow bars in* **Fig. 2a,b**), reflects the sensitivity of the gene knockout to mitotic stopwatch function. As expected, essential genes whose knockout is expected to prolong mitosis, like *MAD2L1BP* and *PLK4*, displayed large mean LFC differences, whereas other essential genes like *GAPDH* dropped out with similar kinetics resulting in a close-to-zero mean LFC difference, and non-essential genes like *OR11H6* exhibited no dropout (**Fig. 2b**).

**Figure 2:**
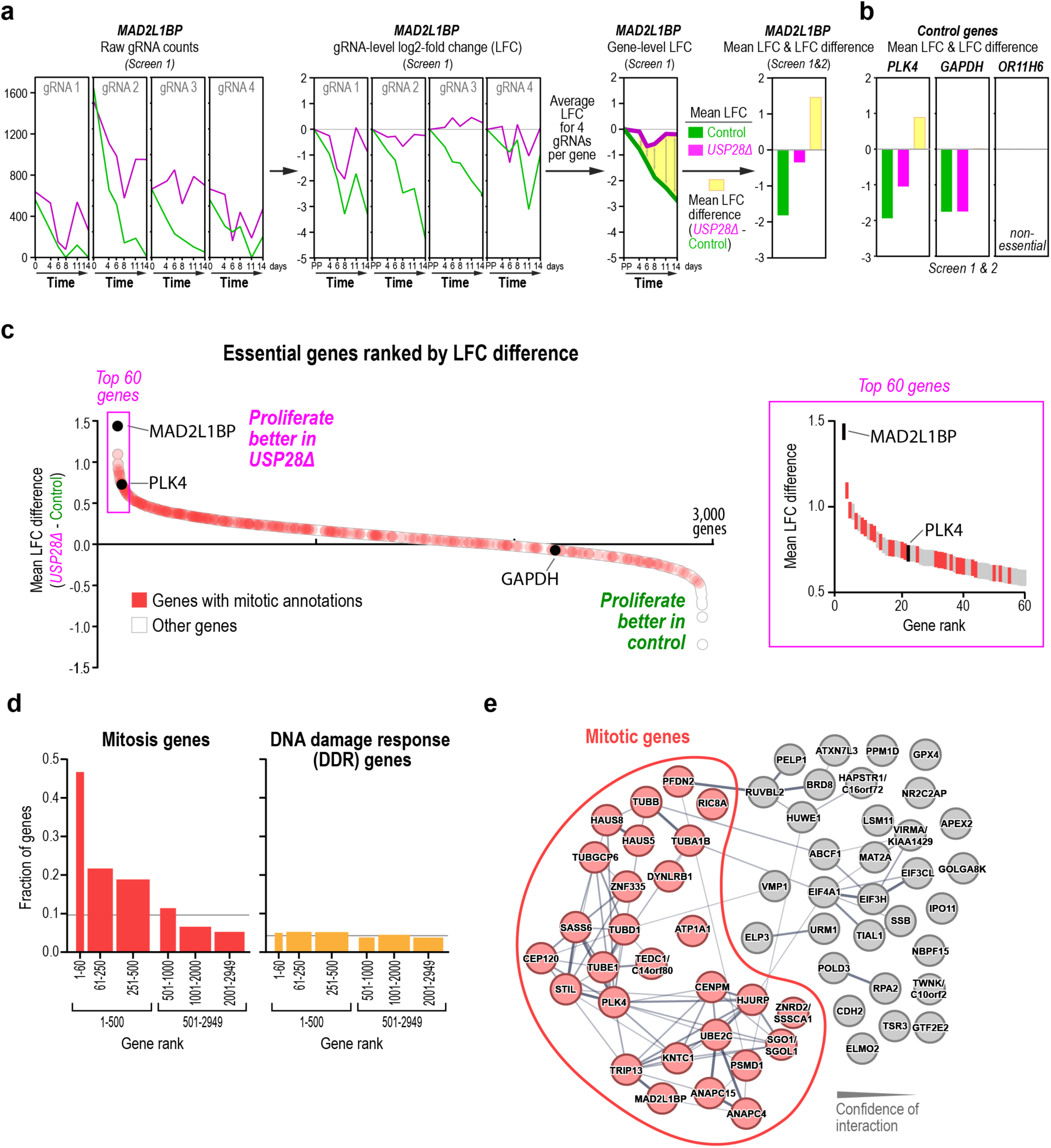
Screening for knockouts sensitive to stopwatch function enriches for mitotic genes. **a,** Example analysis of the screen data for *MAD2L1BP*: (*first panel*) Raw gRNA counts extracted from sequencing data; (*second panel*) LFCs relative to the library plasmid pool (PP) at the assayed time points for each of the 4 gRNAs; (*third panel*) averaged LFCs across all 4 gRNAs; (*fourth panel*) mean LFC across relevant time points averaged between the two screens, and the corresponding difference between the mean LFC in the *USP28Δ* and control cell lines (mean LFC difference). **b,** Screen results for 3 representative genes: the essential mitotic gene *PLK4*, the essential non-mitotic gene *GAPDH*, and *OR11H6*, a non-essential olfactory receptor not expressed in RPE1 cells. **c,** Summary of screen results ranking all 2,953 essential genes by mean LFC difference between the *USP28Δ* and control cell lines, calculated as in *(a)*. Genes with mitotic annotations are highlighted in red. The region containing the 60 genes showing the largest difference between the two cell lines is boxed in magenta and enlarged in the plot on the right. For a summary ranking the 2,953 essential genes by mean LFC difference between the *USP28Δ* and control cell lines see **Supplementary Table 2**. **d,** Graphs highlight the strong enrichment of mitotic (*left*) but not DNA damage response (DDR) genes (*right*) among the top hits. Grey lines indicate the mean fraction of each gene class in the full essential gene set (see also **Extended Data Fig. 1g**). **e,** STRING functional protein association network of the top 60 hits (^28^; line thickness indicates confidence of functional relationship). About half of these hits (*red*) are associated with mitotic annotations and cluster together in the STRING analysis. For nodes where multiple gene names are listed, the first name is the most recent primary gene name, while the second name is how it appears in the screen data.

While the screen was conducted with a genome-wide library, our analysis required gRNA dropout in the control cell line so that we could assess whether or not the dropout rate was impacted by the absence of the stopwatch. To define the set of genes to use for our analysis, we combined a previously identified essential RPE1 gene set^27^ with genes identified as supporting proliferation in our screen, yielding 2,953 genes (**Extended Data Fig. 1**). Ranking these genes by their mean LFC difference revealed that the top 250 genes that exhibited slower gRNA dropout in *USP28Δ* versus control cells, were enriched for mitosis-related functional annotations, whereas the bottom 2,000 genes were depleted for these annotations (**Fig. 2c,d**). In contrast, DNA damage response (DDR) genes showed no enrichment or depletion (**Fig. 2d**; **Extended Data Fig. 1**). STRING network analysis^28^ of the top 60 ranked genes identified a cluster of mitosis-associated genes but also revealed a significant number of genes that lacked a prior direct link to mitotic processes (**Fig. 2e**). We note that the screen also identified genes that exhibited the converse effect – faster gRNA dropout in *USP28Δ* versus control cells. However, as our focus was on knockouts that proliferated better in the absence of the mitotic stopwatch, we did not pursue further analysis of these genes but have provided all of the screen results to support future efforts (Harvard Dataverse repository, https://doi.org/10.7910/DVN/7SFQZ8**; Supplementary Table 2**).

Overall, this unbiased screening approach ranked essential genes based on their sensitivity to the mitotic stopwatch. The top hits of the screen were strongly enriched for, but not exclusively comprised of, mitotic genes.

### A secondary live imaging screen defines two classes of gene knockouts that are sensitive to stopwatch function

The top 60 ranked genes included both genes implicated in mitotic processes and genes that lacked clear association with mitotic events. To determine how many of these 60 genes directly affected mitotic progression, we generated inducible knockout cell lines for each gene in the *USP28Δ* background and performed a live imaging-based secondary screen to quantify mitotic duration following acute (3-day) knockout (**Fig. 3a**; mitotic duration was measured in *USP28Δ* cells to avoid stopwatch-mediated selection prior to imaging). Mitotic events were detected and temporally aligned by chromatin morphology using supervised machine learning with the image analysis software CellCognition^29^ (**Fig. 3a**). To validate this approach, we analyzed acute knockouts of *PLK4* and *MAD2L1BP*, which were both among the top 60 hits. As expected, *PLK4* knockout, which causes centrosome loss and slows spindle assembly^24^, prolonged prometaphase, whereas *MAD2L1BP* knockout, which slows spindle checkpoint silencing^12, 25, 26^, prolonged metaphase (**Fig. 3a**, **Supplementary Movie 1**). Similar analysis of the top 60 gene knockdowns revealed that 27 substantially increased mitotic duration, whereas 33 did not (**Fig. 3b**). We conclude that ranking essential gene knockouts by their ability to proliferate better in the absence of the stopwatch enriches for mitotic genes while also identifying a second distinct class of genes whose acute knockouts do not overtly prolong mitosis.

**Figure 3:**
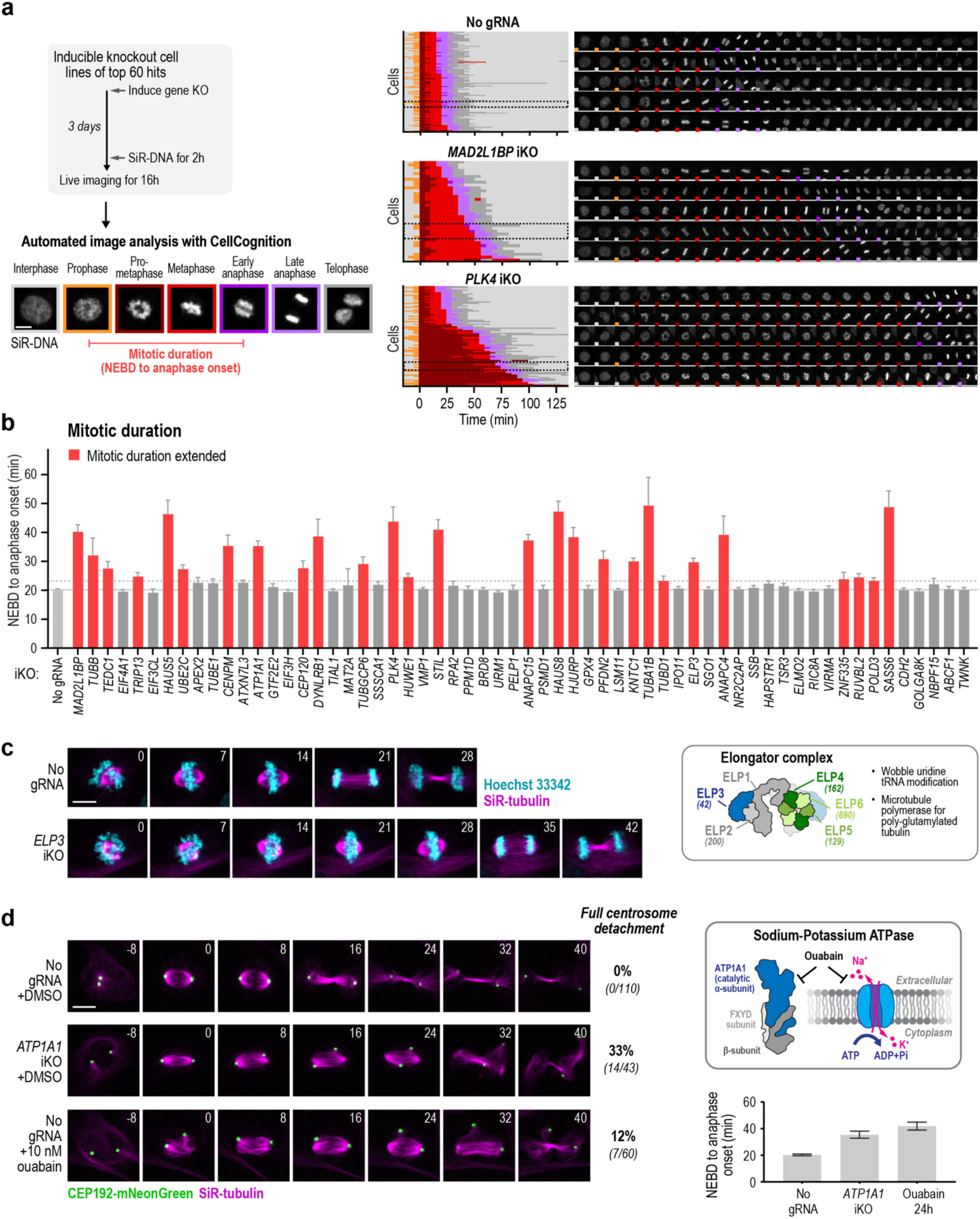
A live imaging secondary screen reveals mitotic phenotypes for knockouts of ∼45% of the top 60 hits. **a,** *Left:* Schematic outline of the method used for the time-lapse imaging secondary screen. Inducible knockout cell lines were generated in the *USP28Δ* background for each of the top 60 hits from the CRISPR screen and cells were imaged 3 days after induction. Movies were analyzed using supervised machine learning in the CellCognition software, which classifies mitotic stages^29^. *Middle:* Each row in the graphs represents a single cell transitioning through mitosis, with stages color-coded as indicated on the left. Individual cells were aligned to the onset of prometaphase, and rows were sorted by the combined duration of prometaphase and metaphase. *Right:* Stills from time-lapse movies corresponding to the boxed regions (*dotted line*) on the graphs. The colored square in the lower left of each still denotes the annotated mitotic stage (see also **Supplementary Movie 1**). **b,** Graph of mean mitotic duration (nuclear envelope breakdown (NEBD) to anaphase onset) for the top 60 hits sorted by screen rank. Error bars represent 95% confidence intervals. The solid line marks the mean in control cells; the dotted line represents the threshold (1.15x of the ‘no gRNA’ control) used to identify genes whose knockout prolongs mitosis. **c,** Stills from time-lapse imaging of chromosomes (*cyan; Hoechst 33342*) and microtubules (*magenta; SiR-tubulin*) after 3-day Cas9 induction in control (WT *USP28*) cells with no gRNA or a gRNA targeting *ELP3* (n=964-1537 cells per condition). Box illustrates the subunit organization of the elongator complex (schematic adapted from ^32^) and its proposed functions. Numbers in brackets below subunit names indicate their ranking in the screen (see also **Supplementary Movie 2**). **d,** *Left:* Stills from live cell imaging monitoring microtubules (*magenta*; SiR-tubulin), and the centrosomal marker CEP192 (*green*) after a 3-day Cas9 induction in control (WT *USP28*) cell lines with no gRNA or a gRNA targeting *ATP1A1*, or after treatment with 10 nM ouabain for 24 h. Frequencies of centrosome detachment from the spindle are indicated (n=43-110 cells per condition). *Right:* Box shows schematic of Na^+^/K^+^–ATPase subunits and its function as a sodium-potassium pump. Graph plots mean mitotic duration (NEBD to anaphase onset) for the indicated conditions (n=133-191 cells per condition) (see also **Extended Data Fig 2 a,b** and **Supplementary Movies 3,4**). Error bars represent 95% confidence intervals. Scale bars, 10 μm.

### Analysis of Class I knockouts reveals a role in mitotic progression for the elongator complex and ATP1A1

The 27 of the top 60 gene hits whose knockouts prolonged mitosis encoded components like PLK4 and MAD2L1BP, loss of which negatively impacts mitosis but does not catastrophically inhibit it. Enrichment for gene knockouts with moderate mitotic defects can be explained by severe mitotic defects resulting in rapid gRNA dropout, limiting the ability to assay for differences in gRNA dropout kinetics between cell lines. Our secondary imaging screen also identified less well-studied genes whose knockout impacted mitotic progression. One such gene is *ELP3*, which encodes a subunit of the elongator complex, a six-subunit complex well-studied for its roles in tRNA modification^30^. Four elongator subunit genes (*ELP2-5*) were among the top 200 screen hits (**Fig. 3c**). Live cell filming revealed that the *ELP3* knockout slowed spindle assembly and chromosome alignment (**Fig. 3c; Supplementary Movie 2**). Recent *in vivo* and *in vitro* biochemical studies have uncovered involvement of the elongator complex in asymmetric spindle assembly and demonstrated its function as a polymerase for polyglutamylated tubulin^31, 32^. Further work will be needed to determine whether elongator involvement in RPE1 cell mitosis depends on its microtubule-directed roles or its canonical function in modifying tRNA wobble-codons to facilitate translation.

A second knockdown that prolonged mitosis was *ATP1A1*, which encodes the catalytic α_1_ subunit of the plasma membrane Na⁺/K⁺-ATPase that employs ATP hydrolysis to export Na⁺ ions and import K⁺ ions, maintaining membrane potential and osmotic balance (^33^; **Fig. 3d**). The pump’s activity is inhibited by ouabain, a cardiac glycoside that binds ATP1A1^34^. *ATP1A1* was also identified in a prior screen profiling essential human genes^35^. To define the consequences of *ATPA1A* loss on mitosis, we imaged RPE1 cells labeled to visualize centrosomes and microtubules. Both *ATP1A1* knockout and low-dose ouabain treatment produced a striking phenotype in which one or both centrosomes were observed to detach from the spindle poles (**Fig. 3d**, **Supplementary Movie 3**); in some cases, centrosomes also shifted or transiently shifted away from the spindle poles without full detachment (**Extended Data Fig. 2,b, Supplementary Movie 4**). We also observed premature (≥ 20 minutes prior to spindle assembly) formation of prominent microtubule asters in ouabain-treated cells (**Extended Data Fig. 2,b**). Collectively, these results suggest that spindle defects contribute to the mitotic delays observed in ATP1A1 knockout cells. Because ouabain inhibits Na⁺/K⁺-ATPase activity without eliminating ATP1A1 protein, these results suggest that the activity of ATP1A1 is important for centrosomes to remain attached to the spindle poles and support normal spindle assembly. These two examples illustrate the ability of the stopwatch screen to identify genes that functionally impact mitotic progression.

### Analysis of Class II knockouts reveals mild p53 and p21 elevation

We next examined the genes identified by our screen that did not appear to prolong mitosis. A STRING network^28^ of the top 60 hits revealed a cluster of genes previously implicated in p53 regulation (**Extended Data Fig. 3**;^36–43^), suggesting that knockdown of these non-mitotic top hits might elevate p53 and its downstream effector p21. To test this, we induced knockout of each non-mitotic top hit for 3 days, fixed the cells, and immunostained for p53 and p21 (**Fig. 4a**). As an internal calibration, we measured responses to Nutlin-3^44^, an MDM2 inhibitor that suppresses p53 ubiquitination (**Fig. 4b**, **Extended Data Fig. 3,c**). For all perturbations, we quantified nuclear p53 and p21 levels using two complementary readouts: (1) the percentage of nuclei with signal above the 98^th^ percentile in DMSO-treated controls (**Fig. 4b,c; Extended Data Fig. 3**) and (2) mean nuclear fluorescence intensity normalized to controls (**Extended Data Fig. 3,d**). Both metrics yielded consistent results. Control and *USP28Δ* cells treated with increasing concentrations (0-8 µM) of MDM2i for 3 or 6 days showed similar p53/p21 induction profiles (**Fig. 4b**, **Extended Data Fig. 3,c**). At low concentration (0.5 µM MDM2i for 3 days), p53 and p21 elevation was just detectable; ∼7-8% of nuclei had p53/p21 levels above the 98^th^ percentile threshold in controls (**Fig. 4b**) and the mean nuclear p53 intensity was ∼1.2-fold that in controls (**Extended Data Fig. 3**). At intermediate concentrations (1-2 µM MDM2i) 20%-60% of cells had p53/p21 levels that exceeded the threshold (**Fig. 4b**) and mean nuclear p53 levels were ∼2-3-fold above controls (**Extended Data Fig. 3**). At the highest MDM2i concentration of 8 µM, nearly all (>90%) cells had p53/p21 levels above the threshold (**Fig. 4b**) and the mean nuclear p53 intensity was ∼6-fold higher than in controls (**Extended Data Fig. 3**). Knockout of each of the 33 non-mitotic top hits elevated either p53 or p21, and all but three elevated both markers (**Fig. 4c,d**). Of the 33 knockouts, 73% caused mild p53/p21 elevation comparable to ∼0.5 µM MDM2i, six produced levels similar to 1 µM MDM2i and three raised p53/p21 levels to an extent between 1 and 2 µM MDM2i (**Fig. 4b,c; Extended Data Fig. 3,c**). Thus, among the top 60 hits, 27 (45%) prolonged mitosis, whereas 33 (55%) elevated p53/p21—generally to a modest extent—without overt mitotic delay (**Fig. 4d**). STRING networks of the top 60 and top 250 hits, highlighting these two gene classes, are shown in *Extended Data Fig. 4*.

**Figure 4:**
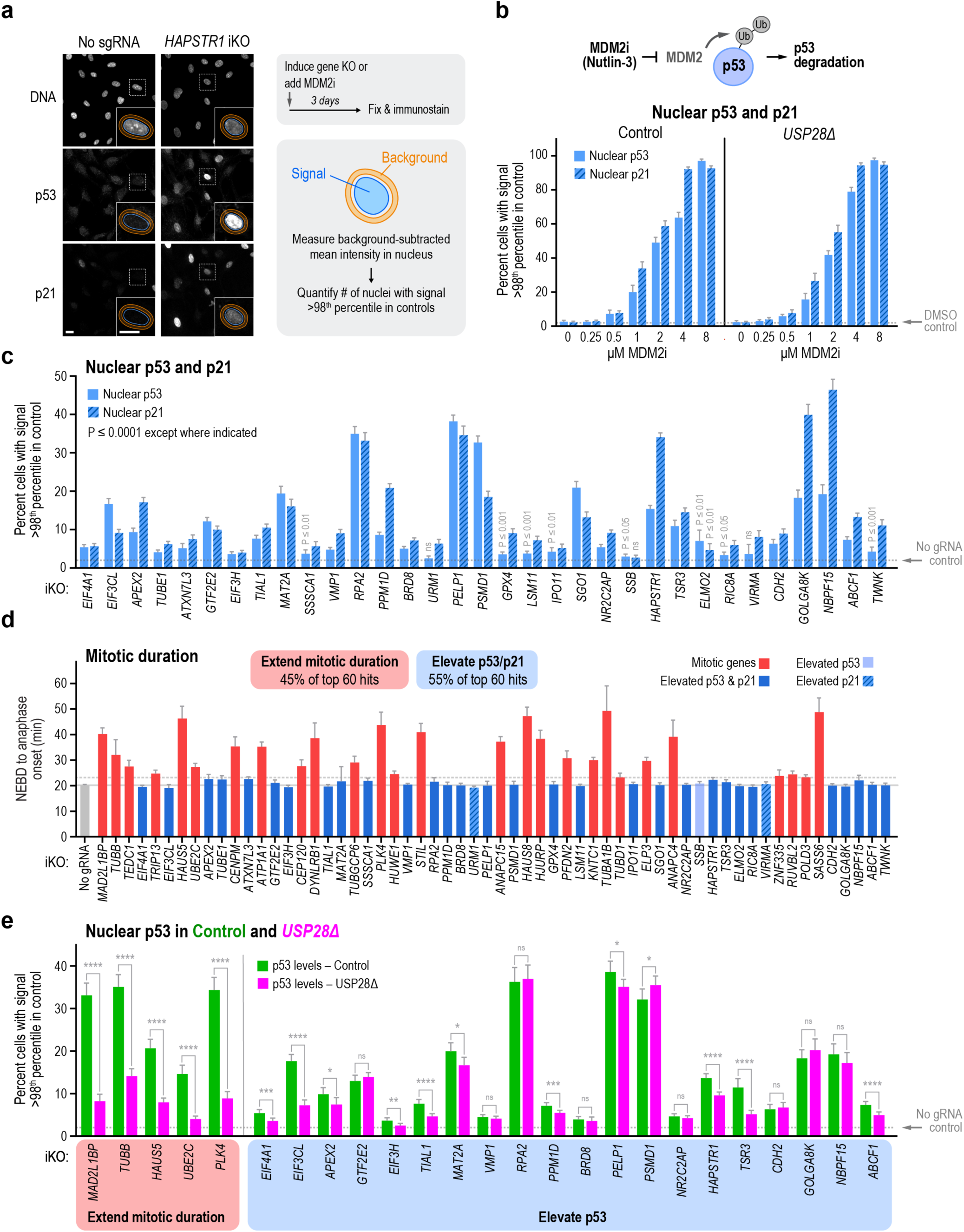
Knockout of non-mitotic top hits leads to mild p53 and p21 elevation. **a,** Immunostaining analysis of nuclear p53 and p21 after inducible knockout of screen hits in control (WT *USP28*) RPE1 cells. Representative images are shown (*left*) together with an overview of the experimental workflow (*right*). Nuclear p53 and p21 intensities were quantified from Hoechst-defined nuclear regions following subtraction of perinuclear background signal. Scale bar, 10 μm. **b,** *Top:* Schematic illustrating MDM2 inhibition by Nutlin-3 (MDM2i). *Bottom:* Percentage of control (*WT USP28*) and *USP28Δ* cells with nuclear p53 (*blue bars*) or p21 (*blue hashed bars*) levels exceeding the 98^th^ percentile of DMSO-treated controls (*dotted line*) after 3-day treatment with the indicated MDM2i concentrations (n=1,517-5,199 cells over 3 replicates per condition). Plots of normalized nuclear fluorescence intensity after 3-day treatment and both measures after 6-day treatment are shown in **Extended Data Fig. 3b,c**. **c,** Quantification of nuclear p53 and p21 levels in control cells 3 days after inducible knockout of top 60 hits that did not prolong mitosis, sorted by screen rank. The graph plots the percentage of cells with nuclear p53 (*blue bars*) or p21 (*blue hashed bars*) above the 98^th^ percentile of ‘no gRNA’ controls (dotted line). Normalized nuclear fluorescence intensity values are shown in **Extended Data Fig. 3d**. Statistical significance was assessed by two-tailed Welch’s t-test. All knockouts differed significantly from the ‘no gRNA’ control (P ≤ 0.0001) unless otherwise indicated (*** = P ≤ 0.001, ** = P ≤ 0.01, * = P ≤ 0.05, ns = P > 0.05; n=1,823-16,158 cells over 3-6 replicates per condition). **d,** Mean mitotic duration for the top 60 hits (*same data as* Fig. 3b), color-coded according to p53/p21 elevation in (*c*). Solid line marks the mean in control cells; dotted line denotes the threshold (1.15x of ‘no gRNA’ control) used to classify knockouts that prolong mitosis. **e,** Comparison of nuclear p53 levels in control (*green bars*) and *USP28Δ* (*magenta bars*) cells 3 days after knockout of genes that prolong mitosis (gene names in *red shaded region*) or genes that elevate p53 levels without prolonging mitosis (gene names in *blue shaded region*). The graph plots the mean percentage of cells exceeding the 98^th^ percentile of ‘no gRNA’ controls (dotted line). Error bars represent 95% confidence intervals. The statistical significance of the difference between the two cell lines was assessed using a two-tailed Welch’s t-test (**** = P ≤ 0.0001, *** = P ≤ 0.001, ** = P ≤ 0.01, * = P ≤ 0.05, ns = P > 0.05; n=1,590-10,979 cells over 3-6 replicates per condition). Normalized nuclear p53 fluorescence intensities, and quantification of p21 using both measures, are shown in **Extended Data Fig. 5a-c**.

To address the contribution of stopwatch activity to p53 elevation, we compared p53 and p21 levels 3 days after knockout of selected genes in the control (intact stopwatch) and *USP28Δ* (no stopwatch) backgrounds. For five knockouts that prolonged mitosis, levels of p53 and p21 were markedly lower in the *USP28Δ* cells compared to controls (**Fig.4e**, **Extended Data Fig. 5-c**), indicating that much of the p53/p21 elevation is a direct consequence of stopwatch activation. By contrast, for the majority of knockouts that elevated p53 without significantly prolonging mitosis, p53 and p21 levels remained similarly high in both backgrounds, consistent with p53 elevation being a stopwatch-independent response to the gene knockout over a three-day interval. Overall, these results support a model in which mild p53 elevation—regardless of its precise cause—synergizes with the mitotic stopwatch to trigger proliferation arrest.

### Mild p53 elevation reduces the mitotic duration threshold of the stopwatch

To determine why knockdowns that modestly elevate p53 without appearing to prolong mitosis render cells sensitive to stopwatch function, we asked whether p53 elevation alters stopwatch behavior. We performed stopwatch assays on cells with mild p53 elevation induced either pharmacologically—with low doses of MDM2 inhibitor (0.25 or 0.5 µM)—or by inducible knockout of two known negative p53 regulators identified in our screen (*PPM1D* and *HAPSTR1*; ^38, 39, 43^). In these assays, asynchronously growing cells are transiently treated with monastrol to generate cells with variable mitotic durations, and the daughters monitored to determine their fate (**Fig. 5a**). In control cells, the majority of daughters of cells that spent longer than 80 minutes in mitosis arrested, whereas those from mothers spending less than 80 minutes in mitosis typically divided again. To test whether pre-existing p53 impacts this temporal threshold, we focused on cells that spent 50-80 minutes in mitosis. 86% of the daughters of DMSO-treated cells with 50-80 min mitoses divided and 14% arrested. When p53 levels were mildly elevated by treatment with 0.25 or 0.5 µM MDM2i, the percentage of daughters that arrested increased to 34 and 52%, respectively (**Fig. 5a,b**). Similarly, while 2.5% of daughters of control cells with no gRNA arrested, 50 and 43% arrested following knockout of *PPM1D* or *HAPSTR1*, respectively (**Fig. 5a,b**). Together, these results support a model in which mild p53 elevation functions as on offset, lowering the amount of mitotic stopwatch complexes that must be produced and inherited to trigger daughter cell arrest (**Fig. 5c**).

**Figure 5:**
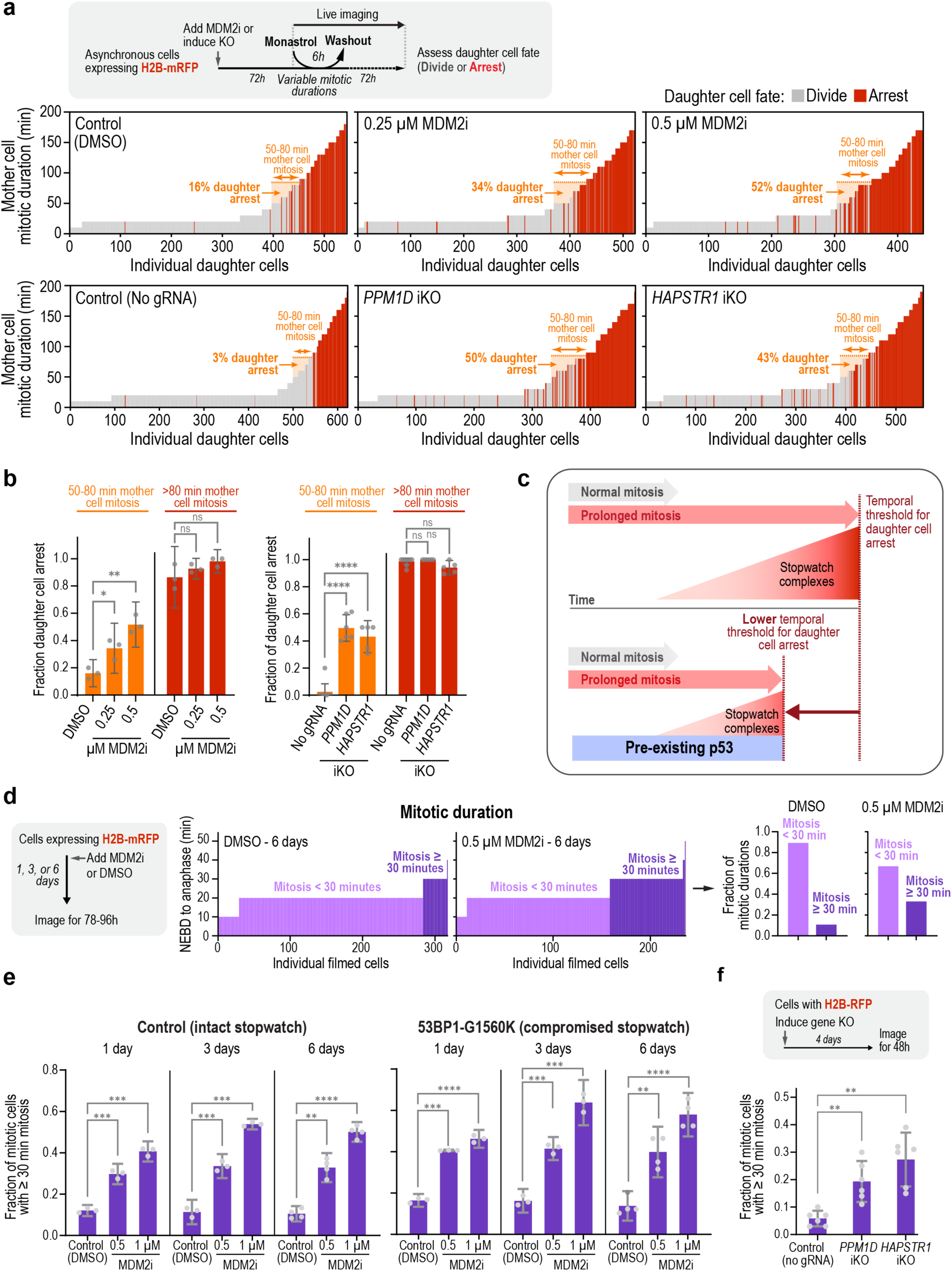
Mild p53 elevation slightly prolongs mitoses and sensitizes the mitotic stopwatch by lowering the temporal threshold for activation. **a,** (*grey box*) Schematic outline of the stopwatch assays performed to assess the relationship between mother cell mitotic duration and daughter cell fate. Assays were performed as in Fig. 1b, except mitotic duration was measured from NEBD to anaphase onset instead of to chromosome decondensation. Stopwatch assays performed in the presence of different doses of MDM2i (*top row*), or after inducible knockout of the negative p53 regulators, *PPM1D* and *HAPSTR1* (*bottom*). The orange shaded region highlights daughter cells whose mothers completed mitosis in 50-80 min; the percentage of these daughters that arrested is indicated. **b,** Mean fraction of daughter cells that arrested after mother cell mitoses >80 min (*red bars*) or between 50-80 min (orange bars) (n=440-622 cells per condition across 3-9 independent experiments). **c,** Schematic illustrating how mild p53 elevation lowers the temporal threshold for daughter cell arrest, thereby sensitizing the mitotic stopwatch. d, *Left:* Schematic of the experiment used to assess impact of persistent mild p53 elevation on mitotic duration. *Middle:* Representative plot of mitotic durations of individual filmed cells. *Right:* Conversion of these data into summary percentages. **e,** Graphs showing the effect of mild persistent p53 elevation on mitotic duration after 1, 3, or 6 days of MDM2i treatment, analyzed as in panel *d*. Both control and a stopwatch-compromised (*TP53BP1-G1560K*) cell line were analyzed (n=174-370 cells per condition across 3 independent experiments). **f,** Mean fraction of cells with mitotic duration ≥ 30 minutes after inducible knockout of the negative p53 regulators, PPM1D and HAPSTR1 for 4 days compared to a ‘no gRNA’ control (n=664-961 cells per condition across 3 independent experiments). Error bars represent 95% confidence intervals. Statistical significance was assessed using a two-tailed Welch’s t-test (**** = P ≤ 0.0001, *** = P ≤ 0.001, ** = P ≤ 0.01, * = P ≤ 0.05, ns = P > 0.05) in all panels.

### Mild p53 elevation subtly extends mitosis

The above results support pre-existing p53 lowering the threshold duration of prolonged mitosis required to arrest daughter cells. However, as normal mitosis is ∼20 min, which is not thought to activate the stopwatch^12^, lowering the temporal threshold for arrest alone cannot explain why knockdowns that modestly elevate p53 proliferate better when the stopwatch is absent. We therefore assessed more carefully whether mild p53 elevation affected mitotic duration. For these experiments, RPE1 cells with an intact mitotic stopwatch were treated with MDM2i for 1, 3, or 6 days and then filmed to monitor mitotic duration (**Fig. 5d,e**). In DMSO-treated controls, ∼90% of mitoses were < 30 minutes and ∼10% were ≥30 minutes. Elevating p53 levels by addition of 0.5 or 1.0 µM MDM2i shifted this distribution, yielding ∼33-50% of mitoses ≥30 minutes. This shift was abolished when p53 was knocked down using an shRNA (**Extended Data Fig. 5**), confirming that it requires p53. Increasing p53 levels by knockout of *PPM1D* or *HAPSTR1* similarly shifted the fraction of mitoses that were ≥30 minutes (**Fig. 5f**). Thus, p53 elevation shifts the distribution of mitotic durations, increasing the frequency of slightly longer (∼30 minute) mitoses. A comparable shift occurred when p53 was elevated by MDM2i treatment in RPE1 cells harboring the 53BP1-G1560K mutation, which compromises the stopwatch by disrupting the interaction between 53BP1 and USP28 (**Fig. 5e**), indicating that this shift to slightly longer mitoses is independent of the stopwatch. We conclude that mild p53 elevation increases the frequency of mitoses with a duration ≥30 minutes. This shift occurs within 1 day of p53 elevation and is independent of stopwatch function (**Fig. 5e**).

### Multi-generational integration enables the stopwatch to detect mild p53 elevation

In prior work, we showed that in RPE1 cells, daughter cells arrest if their mothers undergo a single mitosis longer than ∼90 minutes or two sequential, less prolonged (50-60 minutes) mitoses, indicating that the stopwatch can integrate the effects of extension of mitotic duration over sequential generations^12^. Prolonged mitosis generates stable stopwatch complexes, containing 53BP1 and USP28, that are inherited by daughter cells. This prior work suggested that mild p53 elevation might trigger stopwatch-dependent arrest through the cumulative effect of repeated short mitotic extensions. Consistent with this idea, we noticed that, whereas a gap between *USP28Δ* and control mean LFC plots tended to open up at early timepoints for knockout of mitotic genes, the gap tended to open up later for knockouts whose primary effect was mild p53 elevation (**Extended Data Fig. 5**). This result suggested that while p53/p21 elevation was evident by 3 days after induction of the gene knockouts, the stopwatch-dependent effect on proliferation took additional generations to emerge. To test this idea, we induced p53 elevation by adding 0.5 µM MDM2i, waited 6 days and then performed live cell imaging to record the duration of two successive mitoses (M1 and M2), and followed daughters after the second mitosis to assess their fate (**Fig. 6a**). Each M1–M2 pair was classified as Normal–Normal if both M1 and M2 were < 30 minutes and Mixed/Long-Long if one or both was ≥ 30 minutes (**Fig. 6a**). In control cells with an intact stopwatch, daughters derived from an M1–M2 pair that was Mixed/Long-Long were more than twice as likely to arrest than daughters resulting from a Normal-Normal M1–M2 pair (**Fig. 6a**). This difference was abrogated in cells carrying the 53BP1-G1560K mutation that compromises the mitotic stopwatch (**Fig. 6a**). These results support the idea that mild p53 elevation leads to stopwatch complex-dependent arrest through the cumulative effect of multiple short mitotic extensions in conjunction with reducing the amount of stopwatch complexes that must be accumulated to arrest proliferation (**Fig. 6b**).

**Figure 6:**
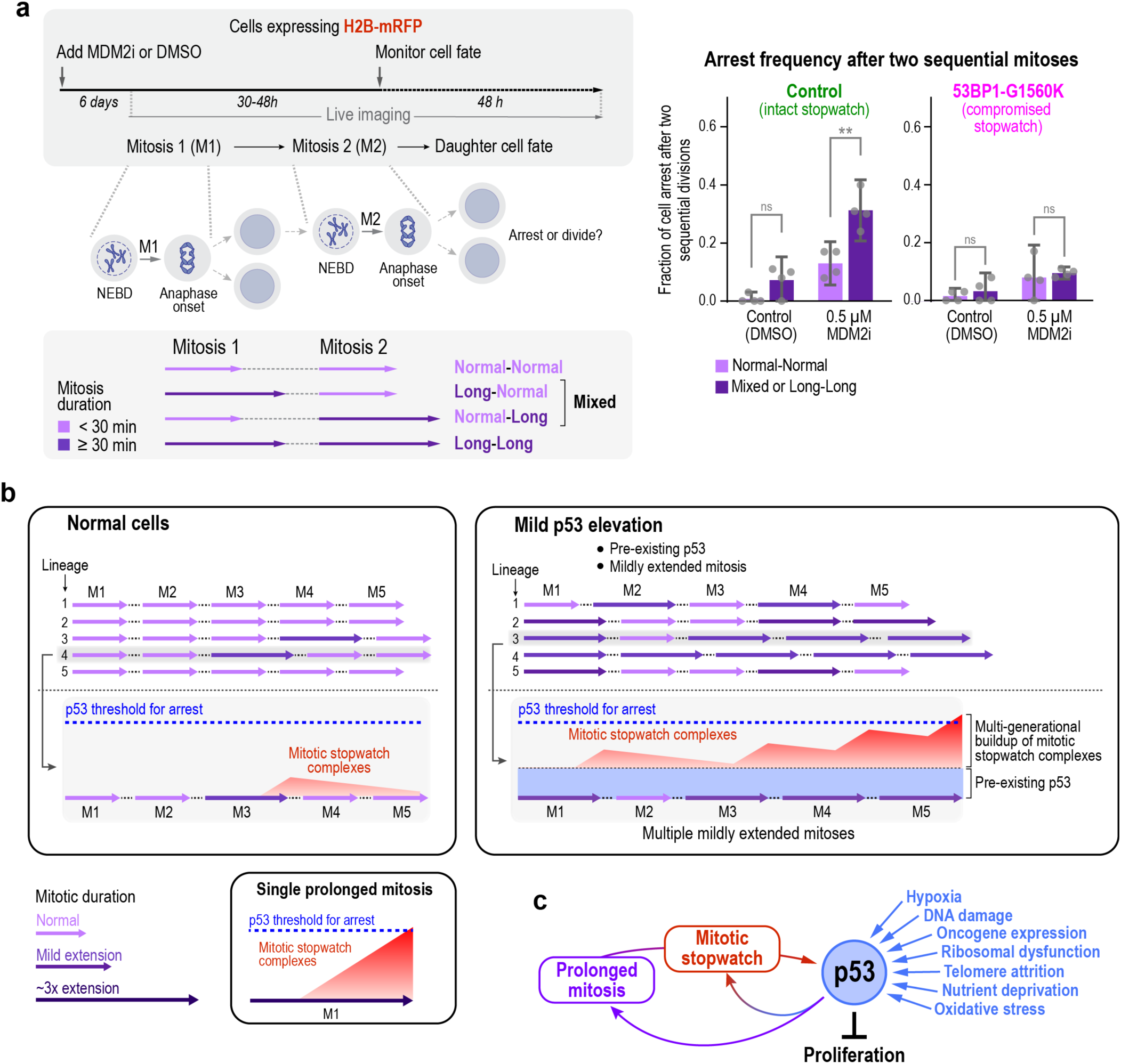
Modest p53 elevation triggers stopwatch-dependent arrest through the cumulative effect of multiple slightly prolonged mitoses. **a,** *Left:* Schematic of experimental approach. Cells were incubated with 0.5 µM MDM2i, or DMSO as a control, for 6 days to induce sustained mild p53 elevation, then filmed for 30-48 h to measure the duration of two sequential mitoses (M1 and M2), followed by 48 h of tracking to assess daughter-cell fate. Each mitosis was classified as < 30 min (*normal*) or ≥ 30 min (*long*), and each M1–M2 pair categorized as Normal–Normal, Mixed, or Long–Long. *Right:* Frequency of daughter cell arrest for the indicated M1–M2 pair classifications in control cells (functional stopwatch) and 53BP1-G1560K cells (compromised stopwatch). Error bars represent 95% confidence intervals; n=56-277 cells per condition across 3 independent experiments. Statistical significance was assessed using two-tailed Welch’s t-test (**** = P ≤ 0.0001, *** = P ≤ 0.001, ** = P ≤ 0.01, * = P ≤ 0.05, ns = P > 0.05). **b,** Model illustrating how the mitotic stopwatch detects mild p53 elevation. *Left:* In cells without elevated p53, slightly prolonged mitoses are uncommon, and the minimal stopwatch complexes they generate dissipate without triggering arrest. *Right:* In cells with modest p53 elevation, pre-existing p53 sensitizes the stopwatch, reducing the amount of stopwatch-derived p53 required for arrest. At the same time, p53 biases cells toward more frequent slightly prolonged mitoses. Because stopwatch complexes are inherited, these incremental events cumulatively increase stopwatch complex levels over successive generations until daughter cell arrest occurs. **c,** Schematic showing how the mitotic stopwatch endows the p53 network with multigenerational memory of diverse stresses. Any cellular stress that causes persistent mild p53 elevation both lowers the threshold for stopwatch-dependent arrest and increases the frequency of slightly prolonged mitoses that generate stopwatch complexes. Together with the strong lineage memory intrinsic to the stopwatch, these effects allow p53 to integrate modest stresses across generations to enforce quality control on cell proliferation.

## Discussion

Recent work has begun to elucidate the mechanism and properties of the mitotic stopwatch, which monitors how long cells spend in mitosis and converts extensions beyond the normal ∼20 minute duration into 53BP1–USP28 complexes that are inherited by daughter cells^9–13^. This inheritance enables the stopwatch to transmit and integrate information across multiple generations. Daughter cells arrest if their mother spends > 90 minutes in mitosis, or if their mother and grandmother each experience moderately prolonged (50-60 minutes) mitoses, indicating that the system possesses a memory that integrates mitotic duration within cell lineages^12^. Because prolonged mitosis is a hallmark of problematic mitoses that can be both symptomatic of and cause chromosome segregation defects, the stopwatch has been proposed to suppress tumorigenesis by preventing the proliferation of potentially dangerous aneuploid progeny^5, 19, 21^. Here, we performed a CRISPR/Cas9 screen comparing gRNA dropout kinetics over multiple generations in RPE1 cells lacking versus possessing the stopwatch to determine whether additional pathways also feed into this signaling system to suppress cell proliferation. By identifying knockouts that dropped out more slowly in cells lacking the stopwatch at any point over a 14-day time course (∼16 generations), we identified essential genes whose loss activated the stopwatch to suppress proliferation.

As expected, genes whose loss is known to directly prolong mitosis—such as *PLK4* and *MAD2L1BP*^12, 24–26^— and genes with mitosis-related annotations were strongly enriched among the top 250 hits (**Fig. 2c,d; Extended Data Fig. 4**). Detailed analysis of the top 60 hits revealed that, in addition to genes whose knockout prolonged mitosis (27 of the top 60 knockouts), there was a second class of genes (33 of the top 60 knockouts) whose knockout did not have an overt effect on mitotic duration (**Figs. 3b, 4d**). Prompted by the presence in this class of a small cluster of known negative regulators of p53 (**Extended Data Fig. 3**), we showed that the primary effect of knockdowns in this class is mild p53 and p21 elevation (**Fig. 4a-d**). Our analysis using both pharmacological and genetic approaches to mildly elevate p53 levels indicates that it has two effects (**Fig. 6b**). First, it sensitizes the mitotic stopwatch: the presence of pre-existing p53 reduces the amount of stopwatch-derived p53 required to trigger arrest. Second, mild p53 elevation shifts the distribution of mitotic timing so that slightly elongated (30 min) mitoses become ∼6-fold more common (∼30%) in cells with elevated p53. Because the stopwatch integrates mitotic duration across generations, the effects of these intermittent slightly prolonged mitoses accumulate to produce sufficient stopwatch complexes and arrest progeny cells (**Fig. 6b**). While our data indicate that mild p53 elevation leads to a slight (∼50%) increase in mitotic duration, the mechanism by which mild p53 elevation prolongs mitosis remains unknown and warrants future investigation.

Overall, our results position the mitotic stopwatch as a central component of the broader p53 stress-response network (**Fig. 6c**). Extensive prior work has shown that p53 is activated by diverse cellular stresses, one of which—prolonged mitosis—is detected through the stopwatch. Our findings suggest that p53 elevation caused by non-mitotic stresses leads to a modest increase in mitotic duration, which is then detected by the stopwatch to arrest proliferation. Because of the memory property of the stopwatch, this slight shift in mitotic timing, combined with stopwatch sensitization by the presence of p53, endows the p53 network with a multigenerational memory of a wide range of stresses. One such stress, as described in a companion paper from another group, is whole genome doubling^45^, but in principle any perturbation that induces mild p53 activation, insufficient on its own to halt proliferation, will engage this mechanism. This ability of the mitotic stopwatch to convert mild, transient mitotic extensions due to low-level p53 elevation into arrest likely explains why its core components, USP28 and 53BP1, have been identified as tumor suppressors and are frequently inactivated in p53-wildtype cancers^16^.

## Methods

### Chemical inhibitors and antibodies

The following chemical inhibitors were used in this study, with their final concentrations, manufacturers, and catalog numbers indicated in parentheses: Monastrol (100 μM; Tocris Biosciences Cat# 1305), Ouabain (10 nM; Sigma Cat# O3125), and Nutlin-3 (MDM2 inhibitor; 0.25-8 μM; MedChemExpress Cat# HY-50696). Antibodies against the following targets are listed here, along with their working concentrations, manufacturers, and catalog numbers: USP28 (1:1,000; Abcam Cat# ab126604, RRID:AB_11127442), p53 (1:100; Millipore Cat# OP43F-100UG, RRID:AB_564964), p21 (1:1,000; Cell Signaling Technology Cat# 2947, RRID:AB_823586), GAPDH (1:1,000; Cell Signaling Technology Cat# 5174, RRID:AB_10622025), p53 (1:1,000; Santa Cruz Biotechnology Cat# sc-126, RRID:AB_628082), p21 (1:1,000; Cell Signaling Technology Cat# 2947, RRID:AB_823586). Secondary antibodies were purchased from GE Healthcare and Jackson ImmunoResearch.

### Cell lines

All cell lines used in this study are listed in **Supplementary Table 1**. RPE1 cell lines were grown in DMEM/F-12 supplemented with 10% FBS, 100 U/ml penicillin, and 100 μg/ml streptomycin at 37°C and 5% CO_2_. The RPE1 control and *USP28Δ* cell lines with inducible Cas9 used for CRISPR screening have been described^12, 24^. Inducible knockout cell lines were generated by lentiviral integration of a gRNA-expressing plasmid based on lentiGuide-Puro^46^ (gift from Feng Zhang; Addgene plasmid # 52963; http://n2t.net/addgene:52963; RRID: Addgene_52963). gRNA sequences were selected based on dropout efficiency in the control cell line in our screens and the Rule Set 2 score for gRNA on-target activity^47^. All gRNA sequences are listed in **Supplementary Table 3**. For transduction of gRNA constructs, harvested virus particles generated as previously described^24^ were added to cells in 6-well plates in the presence of 5 µg/ml polybrene (MilliporeSigma). After 48 h, cells were expanded into 15-cm dishes and selected for 7-10 days in 8 µg/ml puromycin. Cas9 expression was induced with 1 µg/ml doxycycline.

For mitotic stopwatch, mitotic timing, and cell fate assays following sequential mitoses, a plasmid containing H2B-mRFP (EF1α promoter; OD plasmid code pOD3795)^11^ was stably integrated into the genome using lentiviral constructs, and H2B-mRFP-expressing cells were sorted by FACS. For Western blotting, the control and *USP28Δ* iCas9 cell lines were harvested, lysed, and analyzed by immunoblotting as previously described^12^ using the antibodies and concentrations listed above.

### Proliferation assays

For proliferation assays, control or *USP28Δ* iCas9 cells with or without gRNAs for inducible knockouts were plated in triplicate at 50,000 cells per well in a 6-well plate and treated with 1 µg/ml doxycycline to induce Cas9 expression. After 3 days, cells were harvested, counted using a TC20 automated cell counter (Bio-Rad), and re-plated at 50,000 cells per well. Cells were harvested and counted again 6 days after Cas9 induction, and the proliferation from day 3-6 is shown in the graph.

### CRISPR screen

The CRISPR/Cas9 screen was performed using the CRISPR Human Genome 80K Knockout Library in EF1-Puro Vector (Cellecta Cat# KOHGW-EP-80K-V9) containing 4 gRNAs per gene based on the Brunello CRISPR library^47^. For Screen 1, 48 million control or *USP28Δ* RPE1 cells were infected with 48 million infectious virus particles in 14 15-cm plates in 20 ml DMEM/F12 medium containing 8 µg/ml polybrene (MilliporeSigma), corresponding to a multiplicity of infection (MOI) of 1. For Screen 2, half of the quantities were used. After 1 day, 25-27 million cells were transferred to 25-27 15-cm dishes, and infected cells were selected with 10 µg/ml puromycin for 3 days. After 3 days, 25 million cells were seeded in 20 15-cm dishes, and 1 µg/ml doxycycline was added to induce Cas9 expression. Cells were passaged in presence of doxycycline for 15 (Screen 1) or 18 (Screen 2) days, with samples harvested and cells re-seeded on days 2, 4, 6, 8, 11, and 15 post-induction (Screen 1), or on days 2, 4, 7, and 9 for the control and *USP28Δ* cell lines, and on days 12, 15, and 18 for the control cell line (Screen 2). At each time point, samples were harvested by centrifuging trypsinized cells at 277 g for 5 minutes. After removal of the supernatant, cells were washed with 1 ml PBS, and samples were frozen at -80°C for later processing. For Screen 1, harvested samples were isolated and sequenced using the ‘DNA Isolation from Cell Pellets’ and ‘NGS of DNA from Genetic Screen’ services from Cellecta. For Screen 2, DNA was extracted using the QIAamp DNA Blood Maxi Kit (Qiagen) and amplified using the NGS Prep Kit for gRNA Libraries in pRSG16/17 (Cellecta Cat#LNGS-120) along with the Supplementary Primer Set for LNGS-120 (Cellecta Cat# LNGS-120-SP) according to the manufacturer’s protocol. PCR products were gel purified using the QIAquick Gel Extraction Kit (Qiagen) and eluted in 20 µl elution buffer. The gRNA libraries were sequenced on an Illumina NextSeq 500 using gRNA library-specific custom sequencing primers provided in the NGS Prep Kit (Cellecta) according to manufacturer’s instructions. Briefly, primers for sequencing the gRNA sequences (Seq-G, Read 1) and library index sequences (Index-G, Read 2) were spiked into the corresponding Read 1 and Read 2 primer mixes in the NextSeq sequencing reagent cartridge at a final concentration of 300 nM. Libraries were sequenced paired-end for 76 cycles for Read 1 (the gRNA sequence) and 8 cycles for Read 2 (the library index sequence) to a depth of 180 million reads.

### CRISPR screen data analysis

For each screen, raw read counts for each sample were normalized to a fixed total read count of 1 million reads to adjust for variation in sequencing depth, and the control and *USP28Δ* cell lines were analyzed separately. Log2-fold changes (LFCs) at each time point were calculated for each gRNA relative to the read counts in the CRISPR library plasmid pool, which was pre-processed by removing low-count gRNAs with less than 150 reads, and adding a pseudocount of 1 to all reads. Gene-level LFCs were calculated based on the mean of all 4 gRNAs for each time point. To compare differences in dropout kinetics in the presence or absence of the stopwatch, the mean LFC across all time points between day 4 and 14 was calculated for each gene (days 4, 6, 8, 11, and 14 for Screen 1, and days 4, 7, and 9 for Screen 2), and numbers were averaged between both screens. The mean LFC differences between the control and *USP28Δ* cell lines were then calculated by subtracting the mean LFC for the control from the mean LFC for *USP28Δ*, resulting in a single value for each gene. This resulted in positive values for genes whose knockouts drop out more slowly in the absence of the stopwatch. Plots for each gene were generated using R 3.6.1 and deposited in the Harvard Dataverse repository, (https://doi.org/10.7910/DVN/7SFQZ8).

To define a list of essential genes in RPE1 cells, we combined essential genes identified from our data with those from a prior study^27^. In our screens, essential genes were defined as those for which more than half of the gRNAs per gene dropped to LFC ≤ -1.8 between days 6 and 15 in the control cell line in both screens (days 6, 8, 11, and 14 for Screen 1, and days 7, 9, 12, and 15 for Screen 2). The list of 1,432 essential genes that met these criteria was expanded by adding the top 2,500 fitness genes in RPE1 ranked by Bayes Factor in a prior screen^27^, which yielded a total of 2,953 essential genes.

For the gene enrichment analysis of CRISPR screen hits, mitotic genes were defined as those containing mitosis-related keywords in their UniProtKB protein summary (mitosis, spindle, microtubule, tubulin, kinetochore, centromere, centrosome, centriole, cohesion, condensin, dynactin), followed by manual curation to remove genes caught in error by this filter; genes identified as mitotic by a prior screen profiling essential human genes^35^ were also included. DNA damage response (DDR) genes were based on a previously published curated gene list^48^. For STRING^28^ protein association analysis (v12; https://string-db.org), the confidence of functional relationships was determined using data from textmining, experiments, databases, and co-expression.

### High-throughput mitotic duration imaging and data analysis

Cas9 expression in the *USP28Δ* RPE1 cell lines with or without gRNAs for inducible knockouts was induced with 1 µg/ml doxycycline for 3 days before imaging for 48 h. 4,000-5,000 cells per well were seeded into 96-well plates the day before imaging. The DNA marker SiR-DNA (Cytoskeleton Inc) was added 2 h before imaging at a final concentration of 0.5 µM, and cells were imaged for 16 h at 5-minute intervals. Cells were imaged using a CQ1 spinning disk confocal microscope (Yokogawa Electric Corporation) equipped with a 20X 0.8 NA objective, and a 2560x2160 pixel sCMOS camera (Andor) or a 2,000x2,000 pixel sCMOS camera (ORCA-Flash4.0V3, Hamamatsu Photonics) at 37°C and 5% CO_2_. Images were acquired by collecting 5 x 2.5 μm z-stacks using the CQ1 software (Yokogawa).

Mitotic durations were analyzed with the image analysis platform CellCognition (version 1.5.2)^29^, which uses supervised machine learning for classification of cellular morphologies to determine mitotic stages within live-cell imaging data. Mitotic duration was defined as the time from prometaphase to anaphase onset. Data was processed using Python 3.8, and results were manually corrected to remove classification errors due to variable SiR-DNA signal intensities between cell lines.

### Imaging of mitotic phenotypes

For live imaging of DNA and mitotic spindles, control RPE1 cell lines with or without gRNAs were treated with 1 µg/ml doxycycline 3 days before imaging for 48 h to induce Cas9 expression. The day before imaging, 4,000 cells per well were seeded into 96-well plates. The DNA marker Hoechst 33342 (Invitrogen) and microtubule marker SiR-tubulin (Cytoskeleton Inc) were added 2 h before imaging at final concentrations of 50 nM and 20 ng/ml, respectively. Cells were imaged for 24 h collecting 4 x 2.7 μm z-stacks at 7-minute intervals.

For experiments using the Na^+^/K^+^-ATPase inhibitor ouabain, 4,000 *USP28Δ* RPE1 cells, which express *in situ*-tagged CEP192-mNeonGreen, were seeded 2 days before imaging, and 10 nM ouabain was added 24 h before imaging. For mitotic duration analysis, DNA and mitotic spindles were visualized as described above. For imaging CEP192 phenotypes, 8,000 cells were seeded and ouabain was added 24 h before imaging. SiR-tubulin (Cytoskeleton Inc) was added 2-4 h before imaging at a final concentration of 50 nM to visualize microtubules. Cells were imaged for 14-24 h collecting 5 x 2.8 μm z-stacks at 5-8-minute intervals.

### Immunostainings and quantification

For p53 and p21 nuclear quantification, control or *USP28Δ* RPE1 cell lines with inducible Cas9 with or without gRNAs were treated with 1 µg/ml doxycycline 3 days before fixation for 48 h. For MDM2 inhibition experiments, Nutlin-3 was added 1, 3, or 6 days prior to fixation. 3,000-5,000 cells per well were seeded into 96-well plates the day before fixation, and cells were fixed in 100 μl ice-cold methanol for 7 minutes at -20°C and washed twice with washing buffer (PBS containing 0.1% Triton X-100). Cells were blocked with blocking buffer (PBS containing 4% BSA, 0.1% Triton X-100, and 0.05% sodium azide) overnight at 4°C, either on the same day or after storage in PBS at 4°C for 1-3 days. After blocking, cells were incubated with primary antibody in blocking buffer for 2 h (concentrations as indicated above). Cells were washed 3 times with washing buffer and incubated with blocking buffer containing secondary antibodies and 5 μg/mL Hoechst 33342 (Invitrogen) for DNA staining. Finally, cells were washed 3 times with washing buffer before imaging. Imaging was performed on a CQ1 spinning disk confocal system (Yokogawa Electric Corporation) as described above. Images were acquired as 10 x 1 μm z-stacks. Nuclear signal intensities were quantified using the CellPathfinder software (Yokogawa). Nuclear areas were segmented based on DNA staining, along with donut-shaped areas surrounding the nuclei for measuring local background intensities. Background-subtracted p53 and p21 nuclear fluorescence was measured in each nucleus, and results were plotted as the fraction of nuclei with signal intensities above the 98^th^ percentile in ‘no gRNA’ or DMSO controls, or as fluorescence intensities after normalizing controls to 1.

### Mitotic stopwatch, mitotic timing, and cell fate assays following sequential mitoses

For mitotic stopwatch assays, control RPE1 iCas9 cells expressing H2B-mRFP with or without gRNAs for inducible knockouts were either treated with Nutlin-3 for 3 days prior to imaging to inhibit MDM2, or with 1 µg/ml doxycycline 3 days prior to imaging for 48 h to induce Cas9 expression. 1,000-3,000 cells per well were seeded on 96-well plates the day before imaging. 5 x 3 μm z-stacks were acquired on the CQ1 spinning disk confocal system (Yokogawa Electric Corporation) described above, at 10-minute intervals. 100 μM monastrol was added to asynchronously dividing cells just before imaging, and was washed out with pre-equilibrated medium after 6 h, followed by imaging for an additional 72 h. After monastrol washout, cells complete mitosis, and the resulting daughter cells were tracked to determine their fate (arrest or proliferation). Cells that entered mitosis shortly after monastrol washout were also analyzed to represent unperturbed mitotic durations.

For mitotic timing experiments, Nutlin-3 was added 1, 3, or 6 days prior to imaging, or Cas9 expression was induced with 1 µg/ml doxycycline 4 days prior to imaging for 48 h. For cell fate assays following sequential mitoses, cells were treated with Nutlin-3 for 6 days prior to imaging for 78-96 h. The day before imaging, cells were seeded on 96-well plates at 500-1,000 cells per well. Cells were imaged by collecting 5 x 3 μm z-stacks at 10-minute intervals.

### Statistical analysis

P-values were obtained using two-tailed Welch t-tests performed in GraphPad Prism 10. P-values are labelled as follows: **** = P ≤ 0.0001, *** = P ≤ 0.001, ** = P ≤ 0.01, * = P ≤ 0.05, ns = P > 0.05.

### Data availability statement

All data supporting the findings of this study are available within the paper and its Supplementary Information or from the corresponding author upon reasonable request. The complete CRISPR/Cas9 dropout screen dataset, including raw and normalized read counts, log2-fold change values and dropout trajectories (for each gRNA and mean for all 4 gRNAs) for both screens is available at the Harvard Dataverse repository (https://doi.org/10.7910/DVN/7sFQZ8). All RPE1-derived cell lines generated in this study are listed in **Supplementary Table 1** and are available from the corresponding author upon reasonable request, subject to a material transfer agreement where required.

## Acknowledgements

We thank Sven Heinz and Max Chang for sequencing samples from Screen 2, and Mariya Kazachkova for help with analysis of sequencing data. This work was supported by grants from the NIH to A.D. (R01 GM074215) and K.O. (R01 GM074207). B.E.M. was supported by an EMBO Postdoctoral Long-Term Fellowship and an HFSP Postdoctoral Long-Term Fellowship. K.O. acknowledges partial salary support from the Ludwig Institute for Cancer Research. The funders had no role in study design, data collection and analysis, decision to publish, or preparation of the manuscript.

## Author Contributions

Conceptualization: BEM, FM, AD, KO; Methodology: BEM, FM; Formal analysis: BEM, FM; Investigation: BEM, FM; Resources: AD, KO; Data curation: BEM; Writing - original draft: BEM, AD, KO; Writing – review & editing: BEM, FM, AD, KO; Visualization: BEM, AD, KO; Supervision: AD, KO; Project administration: AD, KO; Funding acquisition: AD, KO

## Competing Interests

The authors declare no competing interests.

## Extended Data Figures and Legends

**Extended Data Figure 1:**
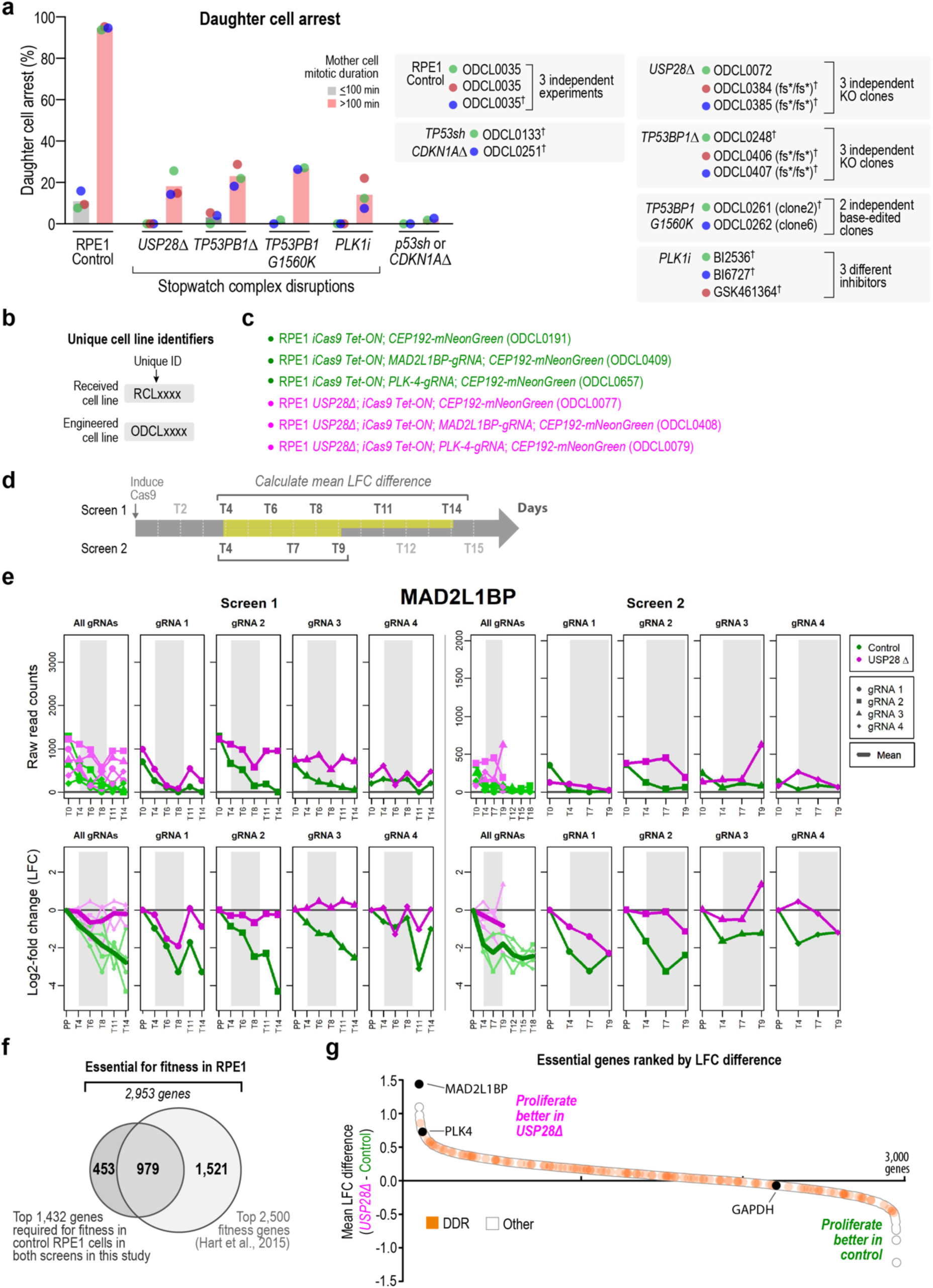
Meta-analysis of prior stopwatch assays, CRISPR screen overview, data display format in accompanying database, and distribution of DNA damage response (DDR) genes across the ranked essential gene set. **a,** Same data as in Fig. 1d except with data points color-coded by experiment. Each stopwatch assay produces a pair of points—corresponding to mother cell durations ≤ 100 min and > 100 min—color-coded by experiment for each condition. Cell line details are provided in the boxes on the right; daggers (†) denote previously published data^12^. **b,** Coding system used to differentiate received and engineered cell lines. See **Supplementary Table 1** for a full list of all cell lines used in this study. **c,** Summary of the features of the RPE1-derived cell lines used in *Fig.1f-h*. **d,** Overview of the CRISPR dropout screen, which was performed twice with samples collected at similar time points. Time points highlighted in bold (range highlighted in yellow) were used to calculate the mean log2-fold change (LFC) difference between the control (*WT USP28*) and *USP28Δ* cell lines, which served as a metric for sensitivity of the gene knockout to stopwatch function. **e,** Example plots for MAD2L1BP from Screen 1 (*left*) and Screen 2 (*right*) formatted as in the dataset accompanying this manuscript (Harvard Dataverse repository, https://doi.org/10.7910/DVN/7SFQZ8). Plots show raw gRNA sequencing counts (*top*) and LFC calculated relative to the library plasmid pool (PP; *bottom*) for each of the 4 gRNAs at the indicated timepoints. In the “All gRNAs” plots, individual traces are shown in lighter colors and the mean across the 4 gRNAs is shown in bold. **f,** A set of 2,953 essential genes in RPE1 cells was compiled by combining 2,500 genes from a previously published dataset^27^ with 453 additional genes identified as essential in both screens performed here. **g,** Same plot as in Fig. 2c, in which all 2,953 essential genes are ranked by mean LFC difference between the *USP28Δ* and control cell lines, except that genes implicated in the DNA damage response (DDR)^48^, rather than genes with mitotic annotations, are highlighted in orange.

**Extended Data Figure 2:**
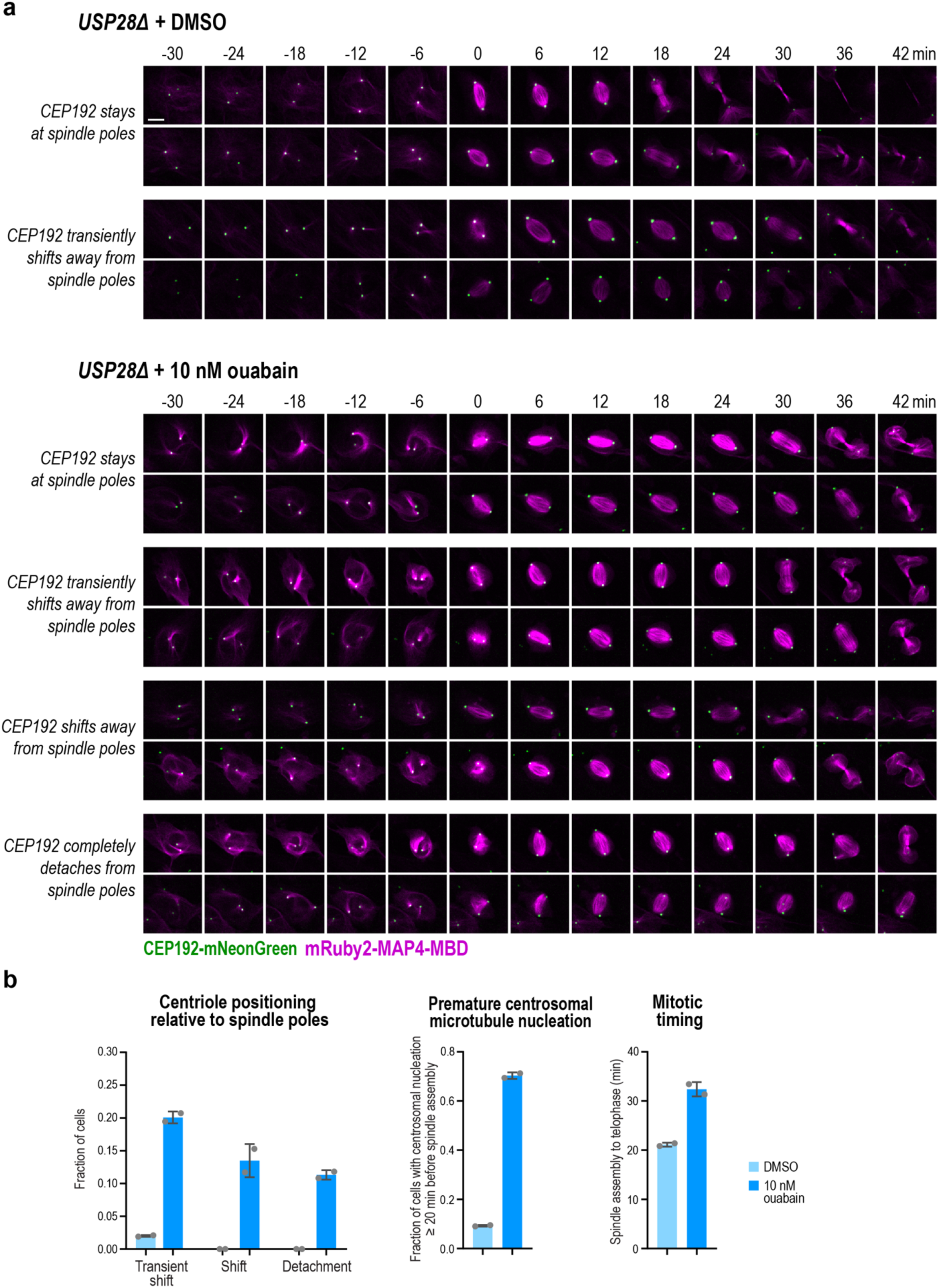
Quantification of mitotic phenotypes resulting from low-dose inhibition of Na⁺/K⁺-ATPase activity. **a,** Stills from time-lapse imaging of control RPE1 cells expressing the centrosomal marker CEP192–mNeonGreen (*green*) and the microtubule marker mRuby2–MAP4-MBD (*magenta; microtubule-binding domain of MAP4 protein*) after treatment with DMSO (*top*) or 10 nM ouabain for 24 h. Representative phenotypes include centrosomes that remain at the spindle poles, detach from the spindle poles, or display transient or sustained shifts away from the poles (see also **Supplementary Movie 4**). Scale bar, 10 μm. **b,** Quantification of the phenotypes shown in (*a*) (*left*), of prominent centrosomal microtubule nucleation occurring ≥ 20 min prior to NEBD (*middle*), and of mitotic duration (spindle assembly to telophase) (*right*) following 24 h treatment with DMSO or 10 nM ouabain. Error bars represent the standard deviation; n=255-301 cells per condition across two replicates.

**Extended Data Figure 3:**
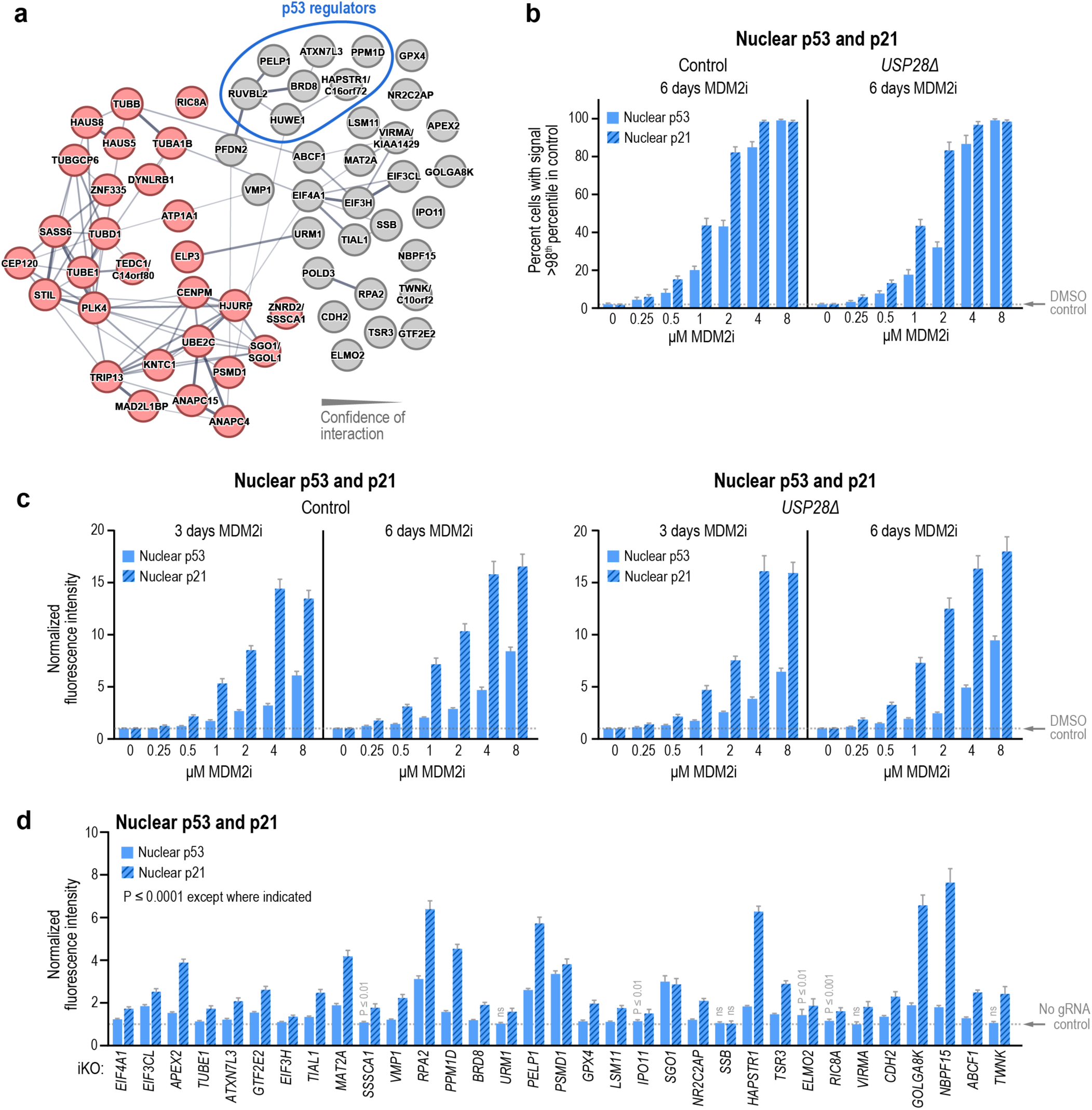
Knockouts of the top hits that do not prolong mitosis elevate p53 and p21. **a,** STRING network^28^ of the top 60 hits reproduced from Fig. 2e highlighting a cluster of genes previously implicated in p53 regulation (*blue circle*). Line thickness indicates confidence of functional interaction. For nodes where multiple gene names are listed, the first name is the most recent primary gene name, while the second name is how it appears in the screen data. **b-c,** Immunostaining was used to analyze nuclear p53 and p21 levels after treatment with increasing concentrations of MDM2i for 3 or 6 days as in Fig. 4a. Graphs plot either the percentage of control (*WT USP28*) and *USP28Δ* cells with nuclear p53 (*blue bars*) or p21 (*blue hashed bars*) above the 98^th^ percentile of DMSO-treated controls (*dotted line*) (*b*) or the normalized mean nuclear fluorescence intensity (*c*) (n=657-5,199 cells over 3 replicates per condition). **d,** Immunostaining was used to analyze nuclear p53 and p21 levels in control cells 3 days after inducible knockout of top 60 hits that did not prolong mitosis, sorted by screen rank. Data correspond to the same experiments used to generate the graph in Fig. 4c, plotted here as mean nuclear fluorescence intensity normalized to the ‘no gRNA’ control (dotted line). Statistical significance was assessed using a two-tailed Welch’s t-test. All knockouts differed significantly from the ‘no gRNA’ control (P ≤ 0.0001) unless otherwise denoted on the graph (*** = P ≤ 0.001, ** = P ≤ 0.01, * = P ≤ 0.05, ns = P > 0.05; n=1,823-16,158 cells over 3-6 replicates per condition). Error bars in all panels represent 95% confidence intervals.

**Extended Data Figure 4:**
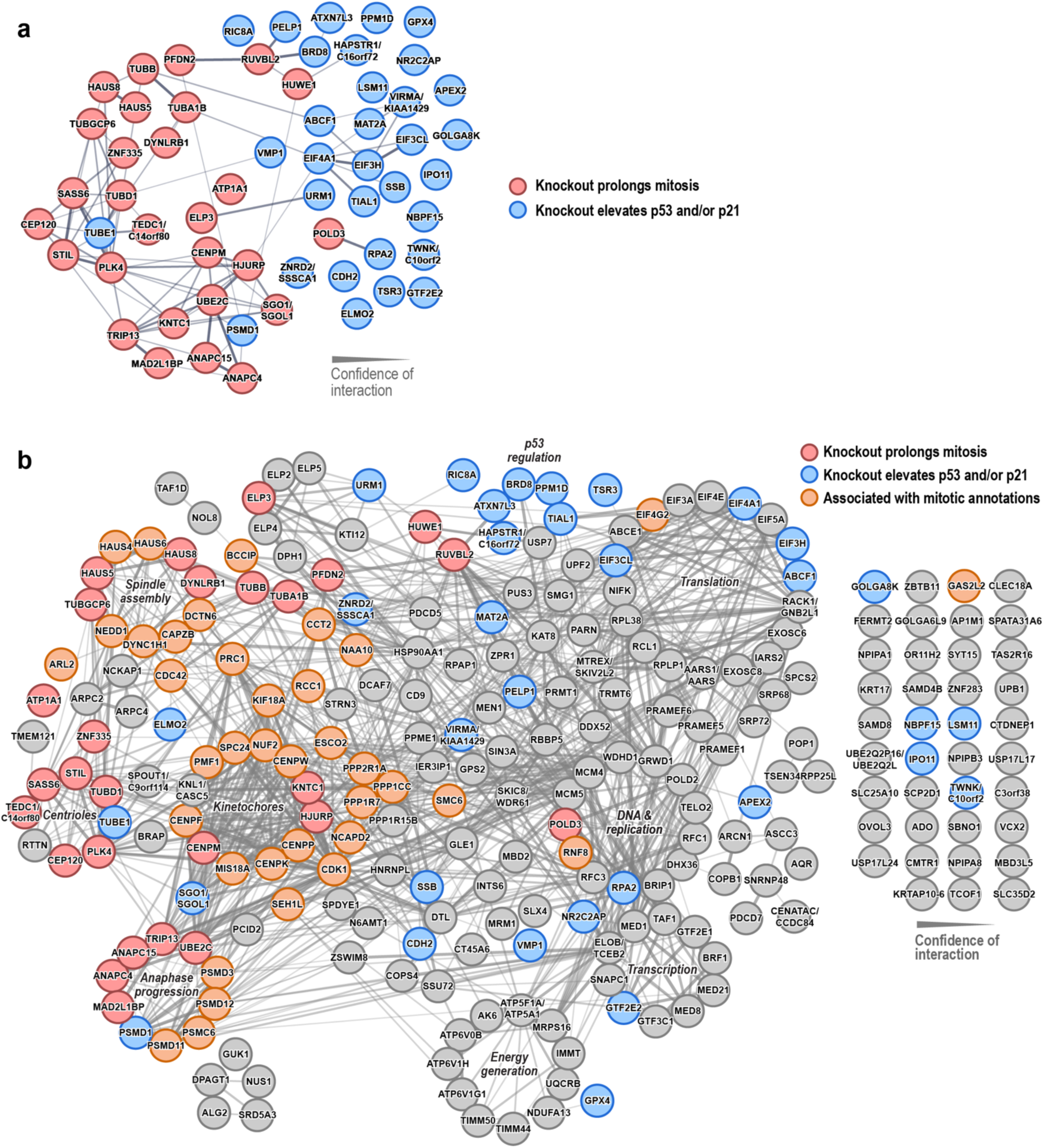
STRING networks of the top 60 and top 250 hits. **a,** STRING network^28^ of the top 60 hits reproduced from Fig. 2e *and Extended Data* Fig. 3a, color-coded based on whether knockout of the gene prolongs mitosis (*red*) or elevates p53/p21 without prolonging mitosis (*blue*) based on data shown in Fig 4d. Line thickness indicates confidence of functional interaction. **b,** STRING network of the top 250 hits, color-coded based on whether their knockout prolongs mitosis (*red*) or leads to p53/p21 elevation without prolonging mitosis (*blue*). Genes ranked beyond the top 60—whose mitotic durations were not directly measured but that are associated with mitotic annotations—are also indicated (*orange*). Line thickness indicates confidence of functional interaction. Labels denote clusters of genes with the indicated functional categories. For nodes where multiple gene names are listed, the first name is the most recent primary gene name, while the second name is how it appears in the screen data.

**Extended Data Figure 5:**
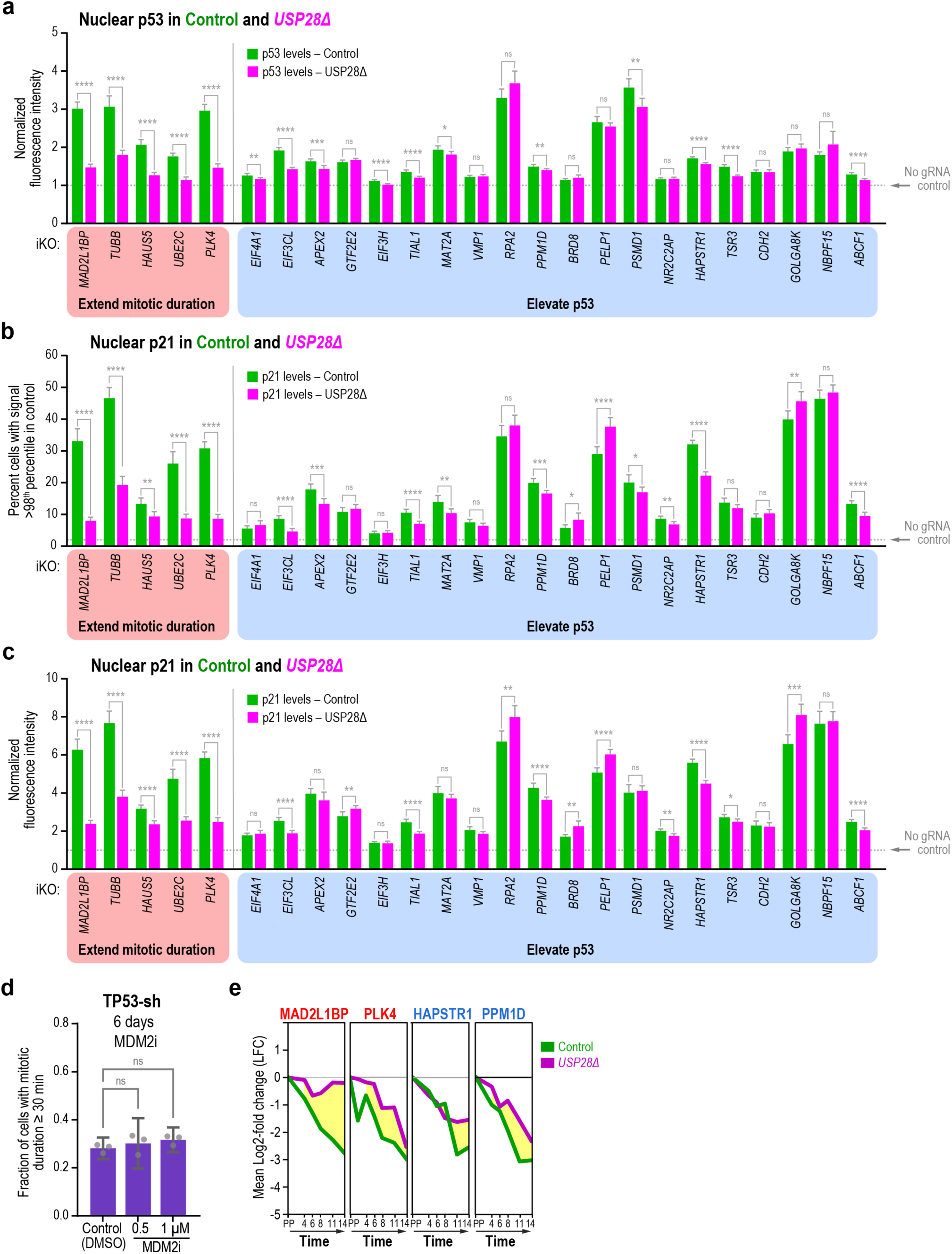
At early timepoints, p53 elevation in knockdowns that prolong mitosis is substantially suppressed by loss of the stopwatch whereas p53 levels in knockdowns whose primary effect is to elevate p53 are not. **a,** Comparison of nuclear p53 levels in control (*green bars*) and *USP28Δ* (magenta bars) cells after knockout of genes that prolong mitosis (*red shaded region*) or genes that elevate p53 levels without prolonging mitosis (*blue shaded region*). Data correspond to the same experiments shown in Fig. 4e, plotted here as mean nuclear fluorescence intensity normalized to the ‘no gRNA’ control (dotted line). Statistical significance between the two cell lines was assessed using a two-tailed Welch’s t-test (**** = P ≤ 0.0001, *** = P ≤ 0.001, ** = P ≤ 0.01, * = P ≤ 0.05, ns = P > 0.05; n=1,590-10,979 cells over 3-6 replicates per condition). **b-c,** Nuclear p21 levels in control (*green bars*) and *USP28Δ* (*magenta bars*) cells were compared after knockout of genes that prolong mitosis (*red shaded region*) or elevate p53 levels without prolonging mitosis (*blue shaded region*). Graphs plot either the percentage of cells with nuclear p21 above the 98^th^ percentile of ‘no gRNA’ controls (*dotted line*) (**b**) or mean nuclear fluorescence intensity normalized to the ‘no gRNA’ control (dotted line) (**c**). Statistical analysis and number of replicates as in (*a*). **d,** Fraction of TP53-sh cells with mitotic duration ≥ 30 minutes after 6 days of MDM2i treatment (n=59-124 cells per condition across 3 independent experiments). Statistical significance was assessed using a two-tailed Welch t-test (**** = P ≤ 0.0001, *** = P ≤ 0.001, ** = P ≤ 0.01, * = P ≤ 0.05, ns = P > 0.05). **e,** CRISPR screen time-course data for the indicated genes. Averaged LFCs relative to the library plasmid pool (PP) across the 4 gRNAs for each gene are plotted versus time for control (*green*) and *USP28Δ* (*magenta*) cell lines. Data are shown for two genes whose knockout prolongs mitosis (*MAD2L1BP* and *PLK4*; red labels) and two negative p53 regulators (*HAPSTR1* and *PPM1D*; blue labels). For *HAPSTR1* and *PPM1D*, the mean LFC difference between control and USP28Δ cells emerges later in the time course than for *MAD2L1BP* and *PLK4*, consistent with a delayed, stopwatch-dependent arrest triggered by modest p53 elevation compared to acute mitotic prolongation. Error bars in *a-d* represent 95% confidence intervals.

## Supplementary Information

**Supplementary Movie 1. Monitoring mitotic duration from time-lapse imaging using automated image analysis.** Spinning-disk confocal imaging using SiR-DNA to mark the chromatin highlighting a metaphase-specific delay in the *MAD2L1BP* knockout and a prometaphase delay in the *PLK4* knockout compared to a ‘no gRNA’ control. Cells were imaged 3 days after knockout induction, and mitotic stages were assigned based on chromatin morphology using the CellCognition software (stage predictions shown as color-coded boxes in the lower-left corner of each frame). Time stamps in the lower right indicate minutes relative to prometaphase onset. Five optical sections (5 x 2.5 μm z-stacks) were acquired every 5 min, and maximal intensity projections were generated for each time point. Playback rate is 1.25 fps. Scale bar, 10 μm. Related to *Fig. 3a*.

**Supplementary Movie 2. Inducible knockout of *ELP3* results in a prolonged metaphase.** Spinning-disk confocal imaging of chromatin labeled with Hoechst 33342 (*cyan*) and microtubules labeled with SiR-tubulin (*magenta*) was used to monitor mitotic spindle dynamics 3 days after inducible knockout of the gene encoding the elongator subunit ELP3. Time stamps in the lower right are minutes after prometaphase onset. 4 optical sections (4 x 2.7 μm) were acquired every 7 min, and maximal intensity projections were generated for each time point. Playback rate is 1.25 fps. Scale bar, 10 μm. Related to *Fig. 3c*.

**Supplementary Movie 3. Knockdown of ATP1A1 or treatment with ouabain reveals detachment of centrosomes from spindle poles and mitotic delay.** Spinning-disk confocal imaging of microtubules labeled with SiR-tubulin (*magenta*) and the stably expressed, *in situ*-tagged centrosomal marker CEP192 (*green*) was performed after either 3-day inducible knockout of ATP1A1 or low-dose treatment with the Na⁺/K⁺-ATPase inhibitor ouabain for 24 h in *USP28Δ* cells. Time stamps in the lower right denote minutes relative to spindle assembly. Yellow arrowheads highlight centrosomes that detach from spindle poles in selected frames. Five optical sections (5 x 2.8 μm) were acquired every 8 min, and maximal intensity projections were generated from z-sections 2-5 for each time point. Playback rate is 1.25 fps. Scale bar, 10 μm. Related to Fig. 3d.

**Supplementary Movie 4: Representative centrosome-positioning defects after ouabain treatment.** Spinning-disk confocal imaging of microtubules labeled with SiR-tubulin (*magenta*) and the stably expressed, *in situ*-tagged centrosomal marker CEP192 (*green*) was performed after low-dose treatment with the Na⁺/K⁺-ATPase inhibitor ouabain for 24 h in *USP28Δ* cells. Centrosome-positioning defects observed range from transient displacements away from the spindle poles to centrosome detachment. Time stamps in the lower right denote minutes relative to spindle assembly. Yellow arrowheads highlight centrosomes that detach from spindle poles in selected frames. Five optical sections (5 x 2.8 μm) were acquired every 6 min, and maximal intensity projections were generated from z-sections 2-5 for each time point. Playback rate is 1.25 fps. Scale bar, 10 μm. Related to *Extended Data Fig. 2*.

## Notes

https://doi.org/10.7910/DVN/7SFQZ8

### Competing Interest Statement

The authors have declared no competing interest.

