## Supplementary Table 1 for "The mitotic stopwatch synergizes with mild p53 activation to halt cell proliferation"

**Supplementary Table 1 – Human cell lines used in this study**

**Received cell lines**

| **OD cell line code** | **Parental line** | **Source** | **Clonal or polyclonal** | **Catalog #** |
| --- | --- | --- | --- | --- |
| RCL001 | hTERT  RPE1 | ATCC | N/A | CRL-4000 |

**Engineered cell lines**

| **OD cell line code** | **Parental line** | **Modifications** | **Clonal or polyclonal** | **Reference** |
| --- | --- | --- | --- | --- |
| *Parental cell lines* | | | | |
| ODCL0003 | RCL001 | *CEP192-mNeonGreen* | Clonal | Meitinger et al., 2020 |
| ODCL0059 | RCL001 | *CDKN1A(p21)Δ* | Clonal | Meitinger et al., 2024 |
| ODCL0071 | ODCL0003 | *CEP192-mNeonGreen, USP28Δ* | Clonal | Meitinger et al., 2020 |
| ODCL0247 | RCL001 | *TP53BP1Δ* | Clonal | Meitinger et al., 2016 |
| ODCL0254 | RCL001 | *EF-1⍺pro-APOBEC-1-Cas9(D10A)-UGI; U6pro-gRNA-TP53BP1(G1560K)* | Clone 2 | Meitinger et al., 2024 |
| ODCL0255 | RCL001 | *EF-1⍺pro-APOBEC-1-Cas9(D10A)-UGI; U6pro-gRNA-TP53BP1(G1560K)* | Clone 6 | This study |
| ODCL0381 | RCL001 | *USP28 fs*/fs* (1bp ins)* | Clone 1 | Meitinger et al., 2024 |
| ODCL0382 | RCL001 | *USP28 fs*/fs* (1bp ins)* | Clone 2 | Meitinger et al., 2024 |
| ODCL0395 | ODCL0191 | *CEP192-mNeonGreen,*  *TRE3G^pro^-Cas9, EF-1⍺^pro^-H2B-mRFP* | Polyclonal | This study |
| ODCL0412 | RCL001 | *TP53BP1 fs*/fs* (1bp ins)* | Clone 1 | Meitinger et al., 2024 |
| ODCL0413 | RCL001 | *TP53BP1 fs*/fs* (1bp ins)* | Clone 2 | Meitinger et al., 2024 |
| ODCL0501 | ODCL0191 | *CEP192-mNeonGreen,*  *TRE3G^pro^-Cas9, EF-1⍺^pro^-H2B-mRFP,*  *U6^pro^-HAPSTR1/C16orf72-gRNA* | Polyclonal | This study |
| ODCL0509 | ODCL0191 | *CEP192-mNeonGreen,*  *TRE3G^pro^-Cas9, EF-1⍺^pro^-H2B-mRFP,*  *U6^pro^-PPM1D-gRNA* | Polyclonal | This study |
| *Cell lines used for meta-analysis of prior stopwatch assays* | | | | |
| ODCL0035 | hTERT  RPE1 | *EF-1α^pro^-H2B-mRFP* | Polyclonal | Meitinger et al., 2016 |
| ODCL0072 | ODCL0071 | *CEP192-mNeonGreen, USP28Δ,*  *EF-1α^pro^-H2B-mRFP* | Polyclonal | Meitinger et al., 2020 |
| ODCL0133 | ODCL035 | *TP53-sh,*  *EF-1⍺^pro^-H2B-mRFP* | Polyclonal | Meitinger et al., 2016 |
| ODCL0248 | ODCL0247 | *TP53BP1Δ,*  *EF-1α^pro^-H2B-mRFP* | Polyclonal | Meitinger et al., 2016 |
| ODCL0251 | ODCL059 | *CDKN1A(p21)Δ,*  *EF-1⍺^pro^-H2B-mRFP* | Polyclonal | Meitinger et al., 2024 |
| ODCL0261 | ODCL0254 | *EF-1⍺^pro^-APOBEC-1-Cas9(D10A)-UGI,*  *U6^pro^-gRNA-TP53BP1(G1560K) (clone 2),*  *EF-1⍺^pro^-H2B-mRFP* | Polyclonal | Meitinger et al., 2024 |
| ODCL0262 | ODCL0255 | *EF-1⍺^pro^-APOBEC-1-Cas9(D10A)-UGI,*  *U6^pro^-gRNA-TP53BP1(G1560K) (clone 6),*  *EF-1⍺^pro^-H2B-mRFP* | Polyclonal | Meitinger et al., 2024 |
| ODCL0384 | ODCL0381 | *USP28 fs*/fs* (1bp ins) (clone 1),*  *EF-1α^pro^-H2B-mRFP* | Polyclonal | Meitinger et al., 2024 |
| ODCL0385 | ODCL0382 | *USP28 fs*/fs* (1bp ins) (clone 2),*  *EF-1α^pro^-H2B-mRFP* | Polyclonal | Meitinger et al., 2024 |
| ODCL0406 | ODCL0412 | *TP53BP1 fs*/fs* (1bp ins) (clone 1),*  *EF-1⍺^pro^-H2B-mRFP* | Polyclonal | Meitinger et al., 2024 |
| ODCL0407 | ODCL0413 | *TP53BP1 fs*/fs* (1bp ins) (clone 2),*  *EF-1⍺^pro^-H2B-mRFP* | Polyclonal | Meitinger et al., 2024 |
| *Cell lines used for CRISPR screen* | | | | |
| ODCL0077 | ODCL0071 | *CEP192-mNeonGreen, USP28Δ,*  *TRE3G^pro^-Cas9* | Clonal | Meitinger et al., 2020 |
| ODCL0191 | ODCL0003 | *CEP192-mNeonGreen,*  *TRE3G^pro^-Cas9* | Clonal | Meitinger et al., 2024 |
| *Inducible knockouts of top 60 CRISPR screen hits* | | | | |
| ODCL0079 | ODCL0077 | *CEP192-mNeonGreen, USP28Δ,*  *TRE3G^pro^-Cas9,*  *U6^pro^-PLK4-gRNA* | Polyclonal | Meitinger et al., 2020 |
| ODCL0408 | ODCL0077 | *CEP192-mNeonGreen, USP28Δ,*  *TRE3G^pro^-Cas9,*  *U6^pro^-MAD2L1BP/ p31comet-gRNA* | Polyclonal | Meitinger et al., 2024 |
| ODCL0409 | ODCL0191 | *CEP192-mNeonGreen,*  *TRE3G^pro^-Cas9,*  *U6^pro^-MAD2L1BP/p31comet-gRNA* | Polyclonal | Meitinger et al., 2024 |
| ODCL0637 | ODCL0191 | *CEP192-mNeonGreen,*  *TRE3G^pro^-Cas9,*  *U6^pro^-TUBB-gRNA* | Polyclonal | This study |
| ODCL0638 | ODCL0191 | *CEP192-mNeonGreen,*  *TRE3G^pro^-Cas9,*  *U6^pro^-TEDC1/C14orf80-gRNA* | Polyclonal | This study |
| ODCL0639 | ODCL0191 | *CEP192-mNeonGreen,*  *TRE3G^pro^-Cas9,*  *U6^pro^-EIF4A1-gRNA* | Polyclonal | This study |
| ODCL0640 | ODCL0191 | *CEP192-mNeonGreen,*  *TRE3G^pro^-Cas9,*  *U6^pro^-TRIP13-gRNA* | Polyclonal | This study |
| ODCL0641 | ODCL0191 | *CEP192-mNeonGreen,*  *TRE3G^pro^-Cas9,*  *U6^pro^-EIF3CL-gRNA* | Polyclonal | This study |
| ODCL0642 | ODCL0191 | *CEP192-mNeonGreen,*  *TRE3G^pro^-Cas9,*  *U6^pro^-HAUS5-gRNA* | Polyclonal | This study |
| ODCL0643 | ODCL0191 | *CEP192-mNeonGreen,*  *TRE3G^pro^-Cas9,*  *U6^pro^-UBE2C-gRNA* | Polyclonal | This study |
| ODCL0644 | ODCL0191 | *CEP192-mNeonGreen,*  *TRE3G^pro^-Cas9,*  *U6^pro^-APEX2-gRNA* | Polyclonal | This study |
| ODCL0645 | ODCL0191 | *CEP192-mNeonGreen,*  *TRE3G^pro^-Cas9,*  *U6^pro^-TUBE1-gRNA* | Polyclonal | This study |
| ODCL0646 | ODCL0191 | *CEP192-mNeonGreen,*  *TRE3G^pro^-Cas9,*  *U6^pro^-CENPM-gRNA* | Polyclonal | This study |
| ODCL0647 | ODCL0191 | *CEP192-mNeonGreen,*  *TRE3G^pro^-Cas9,*  *U6^pro^-ATXN7L3-gRNA* | Polyclonal | This study |
| ODCL0648 | ODCL0191 | *CEP192-mNeonGreen,*  *TRE3G^pro^-Cas9,*  *U6^pro^-ATP1A1-gRNA* | Polyclonal | This study |
| ODCL0649 | ODCL0191 | *CEP192-mNeonGreen,*  *TRE3G^pro^-Cas9,*  *U6^pro^-GTF2E2-gRNA* | Polyclonal | This study |
| ODCL0650 | ODCL0191 | *CEP192-mNeonGreen,*  *TRE3G^pro^-Cas9,*  *U6^pro^-EIF3H-gRNA* | Polyclonal | This study |
| ODCL0651 | ODCL0191 | *CEP192-mNeonGreen,*  *TRE3G^pro^-Cas9,*  *U6^pro^-CEP120-gRNA* | Polyclonal | This study |
| ODCL0652 | ODCL0191 | *CEP192-mNeonGreen,*  *TRE3G^pro^-Cas9,*  *U6^pro^-DYNLRB1-gRNA* | Polyclonal | This study |
| ODCL0653 | ODCL0191 | *CEP192-mNeonGreen,*  *TRE3G^pro^-Cas9,*  *U6^pro^-TIAL1-gRNA* | Polyclonal | This study |
| ODCL0654 | ODCL0191 | *CEP192-mNeonGreen,*  *TRE3G^pro^-Cas9,*  *U6^pro^-MAT2A-gRNA* | Polyclonal | This study |
| ODCL0655 | ODCL0191 | *CEP192-mNeonGreen,*  *TRE3G^pro^-Cas9,*  *U6^pro^-TUBGCP6-gRNA* | Polyclonal | This study |
| ODCL0656 | ODCL0191 | *CEP192-mNeonGreen,*  *TRE3G^pro^-Cas9,*  *U6^pro^-ZNRD2/SSSCA1-gRNA* | Polyclonal | This study |
| ODCL0657 | ODCL0191 | *CEP192-mNeonGreen,*  *TRE3G^pro^-Cas9,*  *U6^pro^-PLK4-gRNA* | Polyclonal | This study |
| ODCL0658 | ODCL0191 | *CEP192-mNeonGreen,*  *TRE3G^pro^-Cas9,*  *U6^pro^-HUWE1-gRNA* | Polyclonal | This study |
| ODCL0659 | ODCL0191 | *CEP192-mNeonGreen,*  *TRE3G^pro^-Cas9,*  *U6^pro^-VMP1-gRNA* | Polyclonal | This study |
| ODCL0660 | ODCL0191 | *CEP192-mNeonGreen,*  *TRE3G^pro^-Cas9,*  *U6^pro^-STIL-gRNA* | Polyclonal | This study |
| ODCL0661 | ODCL0191 | *CEP192-mNeonGreen,*  *TRE3G^pro^-Cas9,*  *U6^pro^-RPA2-gRNA* | Polyclonal | This study |
| ODCL0662 | ODCL0191 | *CEP192-mNeonGreen,*  *TRE3G^pro^-Cas9,*  *U6^pro^-PPM1D-gRNA* | Polyclonal | This study |
| ODCL0663 | ODCL0191 | *CEP192-mNeonGreen,*  *TRE3G^pro^-Cas9,*  *U6^pro^-BRD8-gRNA* | Polyclonal | This study |
| ODCL0664 | ODCL0191 | *CEP192-mNeonGreen,*  *TRE3G^pro^-Cas9,*  *U6^pro^-URM1-gRNA* | Polyclonal | This study |
| ODCL0665 | ODCL0191 | *CEP192-mNeonGreen,*  *TRE3G^pro^-Cas9,*  *U6^pro^-PELP1-gRNA* | Polyclonal | This study |
| ODCL0666 | ODCL0191 | *CEP192-mNeonGreen,*  *TRE3G^pro^-Cas9,*  *U6^pro^-ANAPC15-gRNA* | Polyclonal | This study |
| ODCL0667 | ODCL0191 | *CEP192-mNeonGreen,*  *TRE3G^pro^-Cas9,*  *U6^pro^-PSMD1-gRNA* | Polyclonal | This study |
| ODCL0668 | ODCL0191 | *CEP192-mNeonGreen,*  *TRE3G^pro^-Cas9,*  *U6^pro^-HAUS8-gRNA* | Polyclonal | This study |
| ODCL0669 | ODCL0191 | *CEP192-mNeonGreen,*  *TRE3G^pro^-Cas9,*  *U6^pro^-HJURP-gRNA* | Polyclonal | This study |
| ODCL0670 | ODCL0191 | *CEP192-mNeonGreen,*  *TRE3G^pro^-Cas9,*  *U6^pro^-GPX4-gRNA* | Polyclonal | This study |
| ODCL0671 | ODCL0191 | *CEP192-mNeonGreen,*  *TRE3G^pro^-Cas9,*  *U6^pro^-PFDN2-gRNA* | Polyclonal | This study |
| ODCL0672 | ODCL0191 | *CEP192-mNeonGreen,*  *TRE3G^pro^-Cas9,*  *U6^pro^-LSM11-gRNA* | Polyclonal | This study |
| ODCL0673 | ODCL0191 | *CEP192-mNeonGreen,*  *TRE3G^pro^-Cas9,*  *U6^pro^-KNTC1-gRNA* | Polyclonal | This study |
| ODCL0674 | ODCL0191 | *CEP192-mNeonGreen,*  *TRE3G^pro^-Cas9,*  *U6^pro^-TUBA1B-gRNA* | Polyclonal | This study |
| ODCL0675 | ODCL0191 | *CEP192-mNeonGreen,*  *TRE3G^pro^-Cas9,*  *U6^pro^-TUBD1-gRNA* | Polyclonal | This study |
| ODCL0676 | ODCL0191 | *CEP192-mNeonGreen,*  *TRE3G^pro^-Cas9,*  *U6^pro^-IPO11-gRNA* | Polyclonal | This study |
| ODCL0677 | ODCL0191 | *CEP192-mNeonGreen,*  *TRE3G^pro^-Cas9,*  *U6^pro^-ELP3-gRNA* | Polyclonal | This study |
| ODCL0678 | ODCL0191 | *CEP192-mNeonGreen,*  *TRE3G^pro^-Cas9,*  *U6^pro^-SGO1/SGOL1-gRNA* | Polyclonal | This study |
| ODCL0679 | ODCL0191 | *CEP192-mNeonGreen,*  *TRE3G^pro^-Cas9,*  *U6^pro^-ANAPC4-gRNA* | Polyclonal | This study |
| ODCL0680 | ODCL0191 | *CEP192-mNeonGreen,*  *TRE3G^pro^-Cas9,*  *U6^pro^-NR2C2AP-gRNA* | Polyclonal | This study |
| ODCL0681 | ODCL0191 | *CEP192-mNeonGreen,*  *TRE3G^pro^-Cas9,*  *U6^pro^-SSB-gRNA* | Polyclonal | This study |
| ODCL0682 | ODCL0191 | *CEP192-mNeonGreen,*  *TRE3G^pro^-Cas9,*  *U6^pro^-HAPSTR1/C16orf72-gRNA* | Polyclonal | This study |
| ODCL0683 | ODCL0191 | *CEP192-mNeonGreen,*  *TRE3G^pro^-Cas9,*  *U6^pro^-TSR3-gRNA* | Polyclonal | This study |
| ODCL0684 | ODCL0191 | *CEP192-mNeonGreen,*  *TRE3G^pro^-Cas9,*  *U6^pro^-ELMO2-gRNA* | Polyclonal | This study |
| ODCL0685 | ODCL0191 | *CEP192-mNeonGreen,*  *TRE3G^pro^-Cas9,*  *U6^pro^-RIC8A-gRNA* | Polyclonal | This study |
| ODCL0686 | ODCL0191 | *CEP192-mNeonGreen,*  *TRE3G^pro^-Cas9,*  *U6^pro^-VIRMA/KIAA1429-gRNA* | Polyclonal | This study |
| ODCL0687 | ODCL0191 | *CEP192-mNeonGreen,*  *TRE3G^pro^-Cas9,*  *U6^pro^-ZNF335-gRNA* | Polyclonal | This study |
| ODCL0688 | ODCL0191 | *CEP192-mNeonGreen,*  *TRE3G^pro^-Cas9,*  *U6^pro^-RUVBL2-gRNA* | Polyclonal | This study |
| ODCL0689 | ODCL0191 | *CEP192-mNeonGreen,*  *TRE3G^pro^-Cas9,*  *U6^pro^-POLD3-gRNA* | Polyclonal | This study |
| ODCL0690 | ODCL0191 | *CEP192-mNeonGreen,*  *TRE3G^pro^-Cas9,*  *U6^pro^-SASS6-gRNA* | Polyclonal | This study |
| ODCL0691 | ODCL0191 | *CEP192-mNeonGreen,*  *TRE3G^pro^-Cas9,*  *U6^pro^-CDH2-gRNA* | Polyclonal | This study |
| ODCL0692 | ODCL0191 | *CEP192-mNeonGreen,*  *TRE3G^pro^-Cas9,*  *U6^pro^-GOLGA8K-gRNA* | Polyclonal | This study |
| ODCL0693 | ODCL0191 | *CEP192-mNeonGreen,*  *TRE3G^pro^-Cas9,*  *U6^pro^-NBPF15-gRNA* | Polyclonal | This study |
| ODCL0694 | ODCL0191 | *CEP192-mNeonGreen,*  *TRE3G^pro^-Cas9,*  *U6^pro^-ABCF1-gRNA* | Polyclonal | This study |
| ODCL0695 | ODCL0191 | *CEP192-mNeonGreen,*  *TRE3G^pro^-Cas9,*  *U6^pro^-TWNK/C10orf2-gRNA* | Polyclonal | This study |
| ODCL0696 | ODCL0077 | *CEP192-mNeonGreen, USP28Δ,*  *TRE3G^pro^-Cas9,*  *U6^pro^-TUBB-gRNA* | Polyclonal | This study |
| ODCL0697 | ODCL0077 | *CEP192-mNeonGreen, USP28Δ,*  *TRE3G^pro^-Cas9,*  *U6^pro^-TEDC1/C14orf80-gRNA* | Polyclonal | This study |
| ODCL0698 | ODCL0077 | *CEP192-mNeonGreen, USP28Δ,*  *TRE3G^pro^-Cas9,*  *U6^pro^-EIF4A1-gRNA* | Polyclonal | This study |
| ODCL0699 | ODCL0077 | *CEP192-mNeonGreen, USP28Δ,*  *TRE3G^pro^-Cas9,*  *U6^pro^-TRIP13-gRNA* | Polyclonal | This study |
| ODCL0700 | ODCL0077 | *CEP192-mNeonGreen, USP28Δ,*  *TRE3G^pro^-Cas9,*  *U6^pro^-EIF3CL-gRNA* | Polyclonal | This study |
| ODCL0701 | ODCL0077 | *CEP192-mNeonGreen, USP28Δ,*  *TRE3G^pro^-Cas9,*  *U6^pro^-HAUS5-gRNA* | Polyclonal | This study |
| ODCL0702 | ODCL0077 | *CEP192-mNeonGreen, USP28Δ,*  *TRE3G^pro^-Cas9,*  *U6^pro^-UBE2C-gRNA* | Polyclonal | This study |
| ODCL0703 | ODCL0077 | *CEP192-mNeonGreen, USP28Δ,*  *TRE3G^pro^-Cas9,*  *U6^pro^-APEX2-gRNA* | Polyclonal | This study |
| ODCL0704 | ODCL0077 | *CEP192-mNeonGreen, USP28Δ,*  *TRE3G^pro^-Cas9,*  *U6^pro^-TUBE1-gRNA* | Polyclonal | This study |
| ODCL0705 | ODCL0077 | *CEP192-mNeonGreen, USP28Δ,*  *TRE3G^pro^-Cas9,*  *U6^pro^-CENPM-gRNA* | Polyclonal | This study |
| ODCL0706 | ODCL0077 | *CEP192-mNeonGreen, USP28Δ,*  *TRE3G^pro^-Cas9,*  *U6^pro^-ATXN7L3-gRNA* | Polyclonal | This study |
| ODCL0707 | ODCL0077 | *CEP192-mNeonGreen, USP28Δ,*  *TRE3G^pro^-Cas9,*  *U6^pro^-ATP1A1-gRNA* | Polyclonal | This study |
| ODCL0708 | ODCL0077 | *CEP192-mNeonGreen, USP28Δ,*  *TRE3G^pro^-Cas9,*  *U6^pro^-GTF2E2-gRNA* | Polyclonal | This study |
| ODCL0709 | ODCL0077 | *CEP192-mNeonGreen, USP28Δ,*  *TRE3G^pro^-Cas9,*  *U6^pro^-EIF3H-gRNA* | Polyclonal | This study |
| ODCL0710 | ODCL0077 | *CEP192-mNeonGreen, USP28Δ,*  *TRE3G^pro^-Cas9,*  *U6^pro^-CEP120-gRNA* | Polyclonal | This study |
| ODCL0711 | ODCL0077 | *CEP192-mNeonGreen, USP28Δ,*  *TRE3G^pro^-Cas9,*  *U6^pro^-DYNLRB1-gRNA* | Polyclonal | This study |
| ODCL0712 | ODCL0077 | *CEP192-mNeonGreen, USP28Δ,*  *TRE3G^pro^-Cas9,*  *U6^pro^-TIAL1-gRNA* | Polyclonal | This study |
| ODCL0713 | ODCL0077 | *CEP192-mNeonGreen, USP28Δ,*  *TRE3G^pro^-Cas9,*  *U6^pro^-MAT2A-gRNA* | Polyclonal | This study |
| ODCL0714 | ODCL0077 | *CEP192-mNeonGreen, USP28Δ,*  *TRE3G^pro^-Cas9,*  *U6^pro^-TUBGCP6-gRNA* | Polyclonal | This study |
| ODCL0715 | ODCL0077 | *CEP192-mNeonGreen, USP28Δ,*  *TRE3G^pro^-Cas9,*  *U6^pro^-ZNRD2/SSSCA1-gRNA* | Polyclonal | This study |
| ODCL0716 | ODCL0077 | *CEP192-mNeonGreen, USP28Δ,*  *TRE3G^pro^-Cas9,*  *U6^pro^-HUWE1-gRNA* | Polyclonal | This study |
| ODCL0717 | ODCL0077 | *CEP192-mNeonGreen, USP28Δ,*  *TRE3G^pro^-Cas9,*  *U6^pro^-VMP1-gRNA* | Polyclonal | This study |
| ODCL0718 | ODCL0077 | *CEP192-mNeonGreen, USP28Δ,*  *TRE3G^pro^-Cas9,*  *U6^pro^-STIL-gRNA* | Polyclonal | This study |
| ODCL0719 | ODCL0077 | *CEP192-mNeonGreen, USP28Δ,*  *TRE3G^pro^-Cas9,*  *U6^pro^-RPA2-gRNA* | Polyclonal | This study |
| ODCL0720 | ODCL0077 | *CEP192-mNeonGreen, USP28Δ,*  *TRE3G^pro^-Cas9,*  *U6^pro^-PPM1D-gRNA* | Polyclonal | This study |
| ODCL0721 | ODCL0077 | *CEP192-mNeonGreen, USP28Δ,*  *TRE3G^pro^-Cas9,*  *U6^pro^-BRD8-gRNA* | Polyclonal | This study |
| ODCL0722 | ODCL0077 | *CEP192-mNeonGreen, USP28Δ,*  *TRE3G^pro^-Cas9,*  *U6^pro^-URM1-gRNA* | Polyclonal | This study |
| ODCL0723 | ODCL0077 | *CEP192-mNeonGreen, USP28Δ,*  *TRE3G^pro^-Cas9,*  *U6^pro^-PELP1-gRNA* | Polyclonal | This study |
| ODCL0724 | ODCL0077 | *CEP192-mNeonGreen, USP28Δ,*  *TRE3G^pro^-Cas9,*  *U6^pro^-ANAPC15-gRNA* | Polyclonal | This study |
| ODCL0725 | ODCL0077 | *CEP192-mNeonGreen, USP28Δ,*  *TRE3G^pro^-Cas9,*  *U6^pro^-PSMD1-gRNA* | Polyclonal | This study |
| ODCL0726 | ODCL0077 | *CEP192-mNeonGreen, USP28Δ,*  *TRE3G^pro^-Cas9,*  *U6^pro^-HAUS8-gRNA* | Polyclonal | This study |
| ODCL0727 | ODCL0077 | *CEP192-mNeonGreen, USP28Δ,*  *TRE3G^pro^-Cas9,*  *U6^pro^-HJURP-gRNA* | Polyclonal | This study |
| ODCL0728 | ODCL0077 | *CEP192-mNeonGreen, USP28Δ,*  *TRE3G^pro^-Cas9,*  *U6^pro^-GPX4-gRNA* | Polyclonal | This study |
| ODCL0729 | ODCL0077 | *CEP192-mNeonGreen, USP28Δ,*  *TRE3G^pro^-Cas9,*  *U6^pro^-PFDN2-gRNA* | Polyclonal | This study |
| ODCL0730 | ODCL0077 | *CEP192-mNeonGreen, USP28Δ,*  *TRE3G^pro^-Cas9,*  *U6^pro^-LSM11-gRNA* | Polyclonal | This study |
| ODCL0731 | ODCL0077 | *CEP192-mNeonGreen, USP28Δ,*  *TRE3G^pro^-Cas9,*  *U6^pro^-KNTC1-gRNA* | Polyclonal | This study |
| ODCL0732 | ODCL0077 | *CEP192-mNeonGreen, USP28Δ,*  *TRE3G^pro^-Cas9,*  *U6^pro^-TUBA1B-gRNA* | Polyclonal | This study |
| ODCL0733 | ODCL0077 | *CEP192-mNeonGreen, USP28Δ,*  *TRE3G^pro^-Cas9,*  *U6^pro^-TUBD1-gRNA* | Polyclonal | This study |
| ODCL0735 | ODCL0077 | *CEP192-mNeonGreen, USP28Δ,*  *TRE3G^pro^-Cas9,*  *U6^pro^-IPO11-gRNA* | Polyclonal | This study |
| ODCL0736 | ODCL0077 | *CEP192-mNeonGreen, USP28Δ,*  *TRE3G^pro^-Cas9,*  *U6^pro^-ELP3-gRNA* | Polyclonal | This study |
| ODCL0737 | ODCL0077 | *CEP192-mNeonGreen, USP28Δ,*  *TRE3G^pro^-Cas9,*  *U6^pro^-SGO1/SGOL1-gRNA* | Polyclonal | This study |
| ODCL0738 | ODCL0077 | *CEP192-mNeonGreen, USP28Δ,*  *TRE3G^pro^-Cas9,*  *U6^pro^-ANAPC4-gRNA* | Polyclonal | This study |
| ODCL0739 | ODCL0077 | *CEP192-mNeonGreen, USP28Δ,*  *TRE3G^pro^-Cas9,*  *U6^pro^-NR2C2AP-gRNA* | Polyclonal | This study |
| ODCL0740 | ODCL0077 | *CEP192-mNeonGreen, USP28Δ,*  *TRE3G^pro^-Cas9,*  *U6^pro^-SSB-gRNA* | Polyclonal | This study |
| ODCL0741 | ODCL0077 | *CEP192-mNeonGreen, USP28Δ,*  *TRE3G^pro^-Cas9,*  *U6^pro^-HAPSTR1/C16orf72-gRNA* | Polyclonal | This study |
| ODCL0742 | ODCL0077 | *CEP192-mNeonGreen, USP28Δ,*  *TRE3G^pro^-Cas9,*  *U6^pro^-TSR3-gRNA* | Polyclonal | This study |
| ODCL0743 | ODCL0077 | *CEP192-mNeonGreen, USP28Δ,*  *TRE3G^pro^-Cas9,*  *U6^pro^-ELMO2-gRNA* | Polyclonal | This study |
| ODCL0744 | ODCL0077 | *CEP192-mNeonGreen, USP28Δ,*  *TRE3G^pro^-Cas9,*  *U6^pro^-RIC8A-gRNA* | Polyclonal | This study |
| ODCL0745 | ODCL0077 | *CEP192-mNeonGreen, USP28Δ,*  *TRE3G^pro^-Cas9,*  *U6^pro^-VIRMA/KIAA1429-gRNA* | Polyclonal | This study |
| ODCL0746 | ODCL0077 | *CEP192-mNeonGreen, USP28Δ,*  *TRE3G^pro^-Cas9,*  *U6^pro^-ZNF335-gRNA* | Polyclonal | This study |
| ODCL0747 | ODCL0077 | *CEP192-mNeonGreen, USP28Δ,*  *TRE3G^pro^-Cas9,*  *U6^pro^-RUVBL2-gRNA* | Polyclonal | This study |
| ODCL0748 | ODCL0077 | *CEP192-mNeonGreen, USP28Δ,*  *TRE3G^pro^-Cas9,*  *U6^pro^-POLD3-gRNA* | Polyclonal | This study |
| ODCL0749 | ODCL0077 | *CEP192-mNeonGreen, USP28Δ,*  *TRE3G^pro^-Cas9,*  *U6^pro^-SASS6-gRNA* | Polyclonal | This study |
| ODCL0750 | ODCL0077 | *CEP192-mNeonGreen, USP28Δ,*  *TRE3G^pro^-Cas9,*  *U6^pro^-CDH2-gRNA* | Polyclonal | This study |
| ODCL0751 | ODCL0077 | *CEP192-mNeonGreen, USP28Δ,*  *TRE3G^pro^-Cas9,*  *U6^pro^-GOLGA8K-gRNA* | Polyclonal | This study |
| ODCL0752 | ODCL0077 | *CEP192-mNeonGreen, USP28Δ,*  *TRE3G^pro^-Cas9,*  *U6^pro^-NBPF15-gRNA* | Polyclonal | This study |
| ODCL0753 | ODCL0077 | *CEP192-mNeonGreen, USP28Δ,*  *TRE3G^pro^-Cas9,*  *U6^pro^-ABCF1-gRNA* | Polyclonal | This study |
| ODCL0754 | ODCL0077 | *CEP192-mNeonGreen, USP28Δ,*  *TRE3G^pro^-Cas9,*  *U6^pro^-TWNK/C10orf2-gRNA* | Polyclonal | This study |
| *Cell lines used for imaging mitotic phenotypes or duration* | | | | |
| ODCL0075 | ODCL0071 | *CEP192-mNeonGreen, USP28Δ,*  *EF-1⍺^pro^-mRuby2-MAP4-MBD* | Polyclonal | Meitinger et al., 2020 |
| ODCL0634 | ODCL0395 | *CEP192-mNeonGreen,*  *TRE3G^pro^-Cas9, EF-1⍺^pro^-H2B-mRFP*  *(Generated by FACS sorting the parental cell line for H2B-mRFP expression)* | Polyclonal | This study |
| ODCL0635 | ODCL0501 | *CEP192-mNeonGreen,*  *TRE3G^pro^-Cas9, EF-1⍺^pro^-H2B-mRFP (Generated by FACS sorting the parental cell line for H2B-mRFP expression),*  *U6^pro^-HAPSTR1/C16orf72-gRNA* | Polyclonal | This study |
| ODCL0636 | ODCL0509 | *CEP192-mNeonGreen,*  *TRE3G^pro^-Cas9, EF-1⍺^pro^-H2B-mRFP (Generated by FACS sorting the parental cell line for H2B-mRFP expression),*  *U6^pro^-PPM1D-gRNA* | Polyclonal | This study |

(pro, promoter; ins, insertion)
