## Supplementary Table 3 for "The mitotic stopwatch synergizes with mild p53 activation to halt cell proliferation"

**Supplementary Table 3 – gRNA sequences used for the individual knockouts performed in this study**

| **Gene target** | **gRNA sequence** | **gRNA number** | **Strand** | **Exon** | **gRNA / plasmid name** | **Gene rank in screen** |
| --- | --- | --- | --- | --- | --- | --- |
| ABCF1 | GCAACACATCAATGTTGGGA | 3 | antisense | 12 | pBM0084 | 59 |
| ANAPC15 | ACAGAGTCTCAGTCACACGA | 1 | antisense | 3 | pMF2406 | 31 |
| ANAPC4 | ACATACTACCTTATTCAGCT | 2 | antisense | 18 | pMF2414 | 44 |
| APEX2 | CATGGATGAGTTTACCCAAG | 4 | sense | 3 | pMF2374 | 9 |
| ATP1A1 | CCTGCCTCTTACCGTGACAG | 2 | antisense | 8 | pMF2401 | 13 |
| ATXN7L3 | CATCGCTCAGGAGATATACG | 3 | sense | 2 | pBM0060 | 12 |
| BRD8 | GATATTGCTGTGTCTTACAC | 3 | sense | 11 | pMF2407 | 28 |
| TEDC1/C14orf80 | GGCACTGGCACAACTACCTG | 2 | sense | 3 | pMF2132 | 3 |
| HAPSTR1/C16orf72 | GCAATAAGGATGTGTTGGCT | 2 | sense | 3 | pMF2128 | 47 |
| CDH2 | GTCATGGCAGTAAACTCTGG | 3 | antisense | 8 | pBM0082 | 56 |
| CENPM | CCCCAGTCTCCAGAACACAG | 2 | sense | 4 | pMF2233 | 11 |
| CEP120 | CAGGATCCAAGGCAAAACAT | 2 | antisense | 4 | pMF2402 | 16 |
| DYNLRB1 | CATGAGGCTGGCATACTGGG | 2 | antisense | 3 | pMF2343 | 17 |
| EIF3CL | GCAGAGCATGGTGGTAGATG | 2 | antisense | 15 | pMF2341 | 6 |
| EIF3H | TCTGATCAATGTCCTAATGT | 3 | sense | 5 | pMF2356 | 15 |
| EIF4A1 | CTGCACCTCAGCACGCACGT | 3 | antisense | 5 | pMF2232 | 4 |
| ELMO2 | ACGCAAAGCCATGTACACAA | 2 | sense | 13 | pMF2357 | 49 |
| ELP3 | TACTTCCAACAATATTTACG | 2 | sense | 7 | pMF2167 | 42 |
| GOLGA8K | CCCTCAGGAGCACCTGGAAG | 2 | sense | 14 | pBM0081 | 57 |
| GPX4 | AGAGATCAAAGAGTTCGCCG | 1 | sense | 4 | pMF2409 | 35 |
| GTF2E2 | GAAACAATGGCTAATGACTG | 3 | sense | 4 | pMF2346 | 14 |
| HAUS5 | GCTGTTCGGACATCACGCTA | 3 | antisense | 8 | pMF2129 | 7 |
| HAUS8 | GCTCCCGCTTCCTCTGAGAG | 3 | antisense | 8 | pMF2175 | 33 |
| HJURP | ACCATCAGTGACCTGTACGC | 1 | sense | 8 | pMF2408 | 34 |
| HUWE1 | ATTCTATGCCACATCCTCCG | 3 | sense | 33 | pMF2115 | 23 |
| IPO11 | CTTCAATGAACCAATAAACC | 1 | sense | 4 | pMF2181 | 41 |
| KNTC1 | AAACATTCGGAACACTATGG | 1 | antisense | 26 | pMF2412 | 38 |
| LSM11 | ACTCCATCGCTGTATCCGTG | 2 | sense | 2 | pMF2411 | 37 |
| MAD2L1BP | AAGTGCTTAAGCTGTTCATA | 1 | antisense | 2 | pMF2105 | 1 |
| MAT2A | GTGCAGTATATGCAGGATCG | 3 | sense | 6 | pMF2403 | 19 |
| NBPF15 | GTGGGATCAAGTGAAAAAGG | 2 | sense | 12 | pBM0083 | 58 |
| NR2C2AP | TGGGAGACACGGATGAGCTG | 3 | antisense | 3 | pMF2415 | 45 |
| PELP1 | CCAGCGAGAAGATAGCCTTG | 1 | sense | 15 | pMF2121 | 30 |
| PFDN2 | AGTGATCGATACACTGAAGG | 2 | sense | 3 | pMF2410 | 36 |
| PLK4 | ACTGTGTCAGTGTCGAAGGG | 1 | antisense | 5 | pMF2109 | 22 |
| POLD3 | ATGGCTGAGCTATACACTAG | 1 | sense | 2 | pBM0045 | 54 |
| PPM1D | ATAGCTCCACAAATCACCTG | 2 | antisense | 4 | pMF2384 | 27 |
| PSMD1 | TCACTACACCAAACAATGTG | 3 | sense | 5 | pMF2380 | 32 |
| RIC8A | GCTGCTGGCGCACATCGGTG | 3 | antisense | 3 | pMF2417 | 50 |
| RPA2 | GGAAGTGATCAATGCACACA | 2 | sense | 6 | pMF2405 | 26 |
| RUVBL2 | GATGATTGAGTCCCTGACCA | 2 | sense | 7 | pMF2419 | 53 |
| SASS6 | TATTCGTCTGACTGATGACA | 4 | sense | 3 | pMF2208 | 55 |
| SGO1/SGOL1 | TGGCAGAGATTGGCAAACGC | 3 | sense | 2 | pMF2413 | 43 |
| SSB | CTCTTGATGACATAAAAGAA | 3 | sense | 5 | pMF2416 | 46 |
| STIL | TTACATAAGTCCATATGTGA | 3 | antisense | 13 | pMF2212 | 25 |
| TIAL1 | AATGCGATTGTGCATATGGG | 2 | sense | 7 | pMF2234 | 18 |
| TRIP13 | CATAATTGCAGCAAATCACT | 1 | sense | 3 | pMF2119 | 5 |
| TSR3 | AGGGCCGGCCATAGTTCACG | 1 | antisense | 3 | pBM0061 | 48 |
| TUBA1B | CTGTGATGAGCTGCTCAGGG | 3 | antisense | 3 | pMF2214 | 39 |
| TUBB | CCCCACCGGCACCTACCACG | 2 | sense | 2 | pMF2122 | 2 |
| TUBD1 | ATAGAGGGACAAATACCTAG | 3 | antisense | 5 | pMF2215 | 40 |
| TUBE1 | ACTTTCCAGAATTCACCATG | 2 | antisense | 8 | pMF2239 | 10 |
| TUBGCP6 | GACGCCCCGCTTCACCACAA | 2 | antisense | 4 | pMF2370 | 20 |
| TWNK/C10orf2 | AAATCCGCCAGTATTTGCGG | 1 | sense | 1 | pBM0085 | 60 |
| UBE2C | CTCTCGCTAGAGTTCCCCAG | 2 | sense | 4 | pMF2399 | 8 |
| URM1 | CATGGCTGCGCCCTTGTCAG | 2 | sense | 1 | pMF2379 | 29 |
| VMP1 | AATTCCTAAGCCTATCCAGT | 2 | antisense | 5 | pMF2404 | 24 |
| VIRMA/KIAA1429 | GACATATGATCCATATGACA | 3 | sense | 8 | pMF2184 | 51 |
| ZNF335 | GAAGCCCCACATGTGTGACA | 3 | sense | 12 | pMF2338 | 52 |
| ZNRD2/SSSCA1 | ACCTCGTCCGGAGCACTGTG | 2 | sense | 4 | pMF2345 | 21 |
